# Spinal cord regeneration deploys adult molecular programs that do not recapitulate embryonic development

**DOI:** 10.64898/2026.02.25.708028

**Authors:** Yuxiao Xu, Amulya Saini, Wenda Zhang, Lili Zhou, Mayssa H. Mokalled

## Abstract

Adult zebrafish reverse paralysis after spinal cord injury. Their regenerative capacity is stem cell-dependent and often attributed to potent progenitors that retain embryonic radial glial features and reenact developmental programs after injury. To explore the extents to which regeneration recapitulates development, we integrated single-cell RNA-sequencing datasets spanning development, adult homeostasis and adult regeneration. We found immune cell maturation and neuronal differentiation extend into the juvenile stages, while only 25% of injury-responsive adult progenitors recapitulate larval progenitor identities. By annotating larval progenitors based on their dorso-ventral identities and chronological age, we inferred the spatio-temporal features of adult *sox2^+^* progenitors. This analysis showed the dorso-ventral progenitor identities that guide spinal cord development are not faithfully maintained in homeostatic or regenerating adults. This study reports a genome-wide, single-cell examination of similarities and differences between development and regeneration and indicates adult tissue regeneration repurposes developmental pathways into newly acquired regenerative functions rather than recapitulating development.

## INTRODUCTION

Adult zebrafish are highly regenerative vertebrates that elicit spontaneous cellular repair and functional recovery within 8 weeks of severe spinal cord injury (SCI) (Becker and Becker, 2008; Burris et al., 2021; Tendolkar and Mokalled, 2025). In regenerative vertebrates including zebrafish, spontaneous neural regeneration is stem cell-dependent, driven by potent adult progenitors that proliferate and differentiate to direct neuronal and glial repair (Reimer et al., 2008; Becker and Becker, 2015; Hui et al., 2015; Lindsey et al., 2018; Klatt Shaw et al., 2021; Saraswathy et al., 2022; Zhou et al., 2023). In contrast to zebrafish, spinal cord (SC) progenitors in adult mammals proliferate and differentiate into astrocytes and oligodendrocytes, but exhibit limited inherent ability to differentiate into neurons *in vivo* (Martens et al., 2002; Horky et al., 2006; Meletis et al., 2008; Barnabe-Heider et al., 2010; Karimi-Abdolrezaee et al., 2012; McDonough and Martinez-Cerdeno, 2012; Sabelström et al., 2013; Su et al., 2014; Paniagua-Torija et al., 2018; Shah et al., 2018; Xue et al., 2022). In the absence of potent endogenous stem cells, exogenous stem cell applications have been increasingly used for SCI therapy (Lu et al., 2014; Ceto et al., 2020; Jagrit et al., 2024). We propose that understanding the molecular and spatiotemporal identities of the endogenous stem cells that direct spontaneous SC repair in regenerative animals will guide protocols to improve stem cell-based SCI therapies.

In adult zebrafish, SC progenitors comprise populations of glial cells that line the central canal and express progenitor markers such as *sox2, gfap*, *hey* or *foxj1* (Ogai et al., 2014; Hui et al., 2015; Johnson et al., 2016; Ribeiro et al., 2017). Zebrafish SC progenitors elicit compartmentalized injury responses during SC regeneration. For instance, progenitors within the progenitor motor neuron (pMN) domain express *olig2* and generate motor neurons after SCI (Reimer et al., 2008). On the other hand, ventral progenitors undergo epithelial-to-mesenchymal transition (EMT) and direct glial bridging (Mokalled et al., 2016; Klatt Shaw et al., 2021; Zhou et al., 2023). SC progenitors are typically referred to as ependymal, ependymo-radial glial, radial glial or glial cells, suggesting a hybrid signature between ependymal and radial glial cells (Kroehne et al., 2011; Ogino et al., 2016; Ribeiro et al., 2017). Yet, how the identities of adult progenitors correlate with the radial glial cells that enact embryonic development remains to be investigated.

It is often assumed that the molecular machinery that directs embryonic SC development is deployed to rebuild damaged adult tissues, as multiple developmental pathways are reactivated and required for SCI recovery (Alper and Dorsky, 2022; Swearer et al., 2025). However, even if progenitor cell identities are intrinsically maintained throughout the lifetime of regenerative animals, the extrinsic cues they receive fundamentally differ between development and regeneration. In a first obstacle unique to tissue repair, SC progenitors respond to various immune cells including neutrophils, microglia and T-cells (Hui et al., 2017; Cavone et al., 2021; de Sena-Tomas et al., 2024; Shaw et al., 2024). During later stages of remodeling, regenerating SCs activate canonical developmental pathways such as Fgf signaling to generate new neurons and glia after SCI (Goldshmit et al., 2012). Whereas Fgf2, Fgf3 and Fgf8 promote neurogenesis of *islet1^+^* motor neurons and neurite outgrowth during development, this role is restricted to Fgf3 during regeneration (Goldshmit et al., 2018). Finally, regeneration does not follow the chronologically orchestrated order of developmental events. Thus, it remains unclear how adult progenitors adapt their potency to integrate the extrinsic, spatiotemporal cues of the adult tissue environment to enact regeneration.

Zebrafish embryogenesis is considered complete by 3 days post-fertilization (dpf), when post-embryonic larvae complete hatching and acquire a protruding mouth structure (Kimmel et al., 1995; Parichy et al., 2009). Following larval and juvenile growth (3 to 90 dpf combined), zebrafish adults (>90 dpf) are defined by their ability to produce viable gametes (Kimmel et al., 1995; Parichy et al., 2009). Due to their robust regenerative capacity, larval and adult zebrafish models are increasingly used to elucidate mechanisms of SC regeneration (Vandestadt et al., 2021; Alper and Dorsky, 2022; Tendolkar and Mokalled, 2025). Larvae offer unique benefits such as optical transparency, higher throughput and mapped neuronal circuitry (Alper and Dorsky, 2022). Importantly, while SC regeneration extends over 8 weeks in adults, larval injuries are typically performed at 3 dpf, and swim function is regained within 2 days of injury or 5 dpf (Alper and Dorsky, 2022). Though key regenerative events including immune and progenitor cell activation, axon regeneration and neurogenesis also occur in larvae, differences in timing and cellular mechanisms distinguish larval and adult regeneration. For instance, larvae develop and regenerate in the absence of mature microglia or an adaptive immune system, which emerge between 15 and 35 dpf (Willett et al., 1997; Davidson and Zon, 2004; Xu et al., 2015). Moreover, larval neural stem cells proliferate more rapidly and exhibit greater plasticity than adult progenitors (Park et al., 2007; Anguita-Salinas et al., 2019). Finally, glial bridging is not required for axon regeneration and swim recovery at early stages of development (Wehner et al., 2017; Walker et al., 2025), but is necessary for successful regeneration in 7 dpf larvae and adults. Thus, while targeted genetic studies have been instrumental to establish the requirement of various developmental pathways during regeneration, a distinction between developmental mechanisms that are faithfully recapitulated in regeneration versus developmental pathways that are deployed to acquire regeneration-specific functions is needed.

This is a comprehensive comparative study that reports shared and unique features of larval development and adult SC regeneration in zebrafish. Single-cell RNA-sequencing datasets encompassing larval SC development, adult homeostasis and adult SCI are used (Sur et al., 2023; Saraswathy et al., 2024). We find the timeline for immune cell maturation extends into the juvenile stages, while neurons are markedly more diverse in adults. Delving into progenitor cell identities, only 25% of injury-responsive *sox2*^+^ cell clusters recapitulate radial glial identities from embryonic development. By annotating larval progenitors based on their dorso-ventral identities and chronological maturation, we infer the spatiotemporal features of adult progenitors. In this analysis, 2 out of 10 adult clusters change their maturation scores during regeneration, whereas the dorso-ventral identities that guide SC development are blurred in adult progenitors. This study reveals significant cell identity differences between larval and adult progenitors and indicates adult SC regeneration does not faithfully recapitulate development.

## RESULTS

### Assembling a single-cell atlas of SC development and adult homeostasis in zebrafish

To compare the molecular identities between larval and adult zebrafish, we performed *in silico* integration of multiple RNA-sequencing datasets. For larvae, we used the single-cell RNA-sequencing (scRNA-seq) dataset Daniocell, which was collected from whole embryos between 14 and 120 hours post-fertilization (hpf) (Sur et al., 2023). To account for the majority of SC cells, clusters that were previously classified as “neural”, “SC” or “glial” in DanioCell were included for subsequent analysis (Sur et al., 2023). For adults, we analyzed a single-nuclear RNA-sequencing (snRNA-seq) atlas of dissected SC tissues from adult zebrafish of ∼2 cm in length and at 120 dpf (Saraswathy et al., 2024). In a first analysis, we integrated data from 1, 3 and 5 dpf embryos/larvae along with uninjured adult SCs, capturing key time points of SC development and adult homeostasis **(Fig. 1A)**. A total of 63,353 cells were analyzed, whereby the combined number of larval cells closely matched the number of adult nuclei (30,372 larval cells and 32,972 adult nuclei) **(Fig. 1B)**. Unsupervised clustering of this integration identified 27 clusters with distinct molecular identities **(Fig. 1C)**. Biological samples were well represented and intermixed within each coarse cluster, suggesting our dataset successfully captured biological variation rather than technical or sampling effects **(Fig. S1A, S1B)**. For cell classification, we cloned the previously reported cell identities for each sample onto our integrated dataset **(Fig. S1C)** (Sur et al., 2023; Saraswathy et al., 2024). Coarse clusters were classified as ependymal/glial (EpGlia), neurons, oligodendrocyte precursor cells and oligodendrocytes (OPC/Oligos), immune, as well as endothelial cells and pericytes (Endo/Peri) **(Fig. 1C)**. Cell type annotations were further validated by cross-referencing the top markers of each cluster with canonical cell markers **(Fig. 1D, Table S1)**. *gfap, elavl3, mbpa* and *cd74a* were respectively expressed in EpGlia, neurons, OPC/Oligos and immune clusters **(Fig. 1E)**. This analysis generated an integrated transcriptional atlas of embryonic, larval and adult SC tissues in zebrafish.

**Figure 1.**
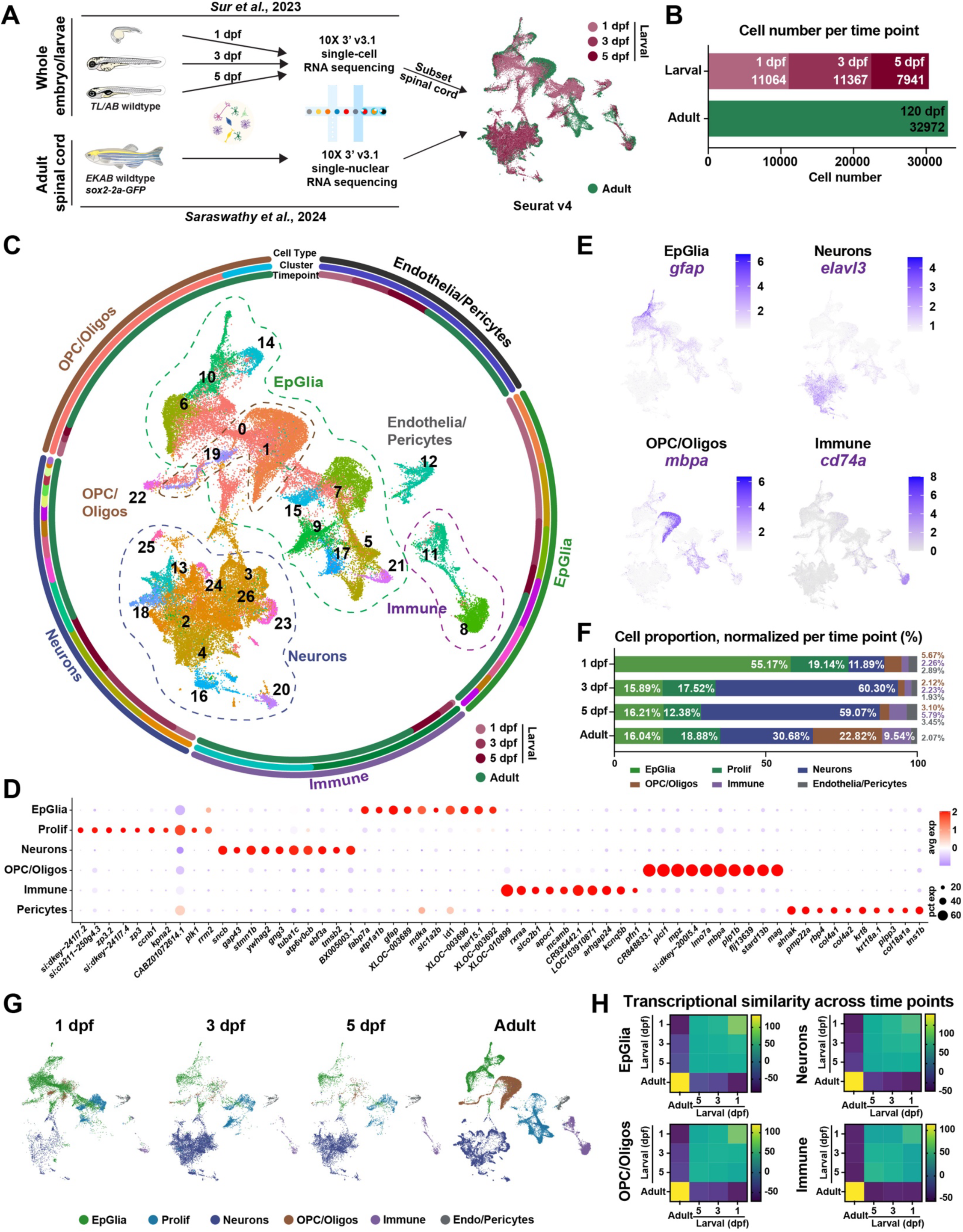
Assembly of a transcriptomic atlas for SC development and adult homeostasis reveals coarse transcriptional differences. **(A)** Single-cell RNA-seq data of zebrafish larval development (DanioCell)(Sur et al., 2023) and single nuclear RNA-seq of adult SC tissues (Saraswathy et al., 2024) were used. For larval analysis, only cells annotated as “neural”, “spinal cord”, “glial” or “immune” from 1, 3 and 5 dpf larvae were included. The adult dataset was generated from dissected SC tissues. **(B)** Numbers of cells analyzed for each time point. The total number of cells from embryo/larvae was comparable to the number of adult cells. **(C)** Circular plot of the larval-adult SC transcriptome. A combined UMAP representation of the integrated dataset shows 27 coarse clusters annotated as EpGlia, neurons, OPC/Oligos and immune cells. Surrounding the UMAP, each circular track presents a metadata column: annotated cell type, cluster number and time point. Cell type and cluster number are annotated and color-coded in the UMAP. Time point colors are listed on the bottom right of the graph. **(D)** Dot plot shows 10 canonical markers for major cell clusters from the integrated dataset. Dot size and color represent percent expressing cells and average gene expression within each cell type, respectively. **(E)** Feature plots show select markers for glia (*gfap*), neurons (*elavl3*), OPC/Oligos (*mbpa*) and immune cells (*cd74a*). **(F)** Cell type composition at 1, 3, 5 and 120 dpf. For each time point and coarse cluster, cell proportions were normalized to the total number of cells from that time point. **(G)** UMAP representation of the integrated larval-adult analysis split by time point. **(H)** ClusterFold similarity analysis across time points. One heatmap is shown for each cell type. Cells from each time point were compared to one another. The extent of transcriptional similarity is represented on a gradient of high (yellow) to low (purple) similarity.

### Adult SC tissues show greater diversity in cell types and molecular profiles

To elucidate the temporal dynamics of SC cells between larval development and adult homeostasis, the numbers and relative proportions of coarse cell clusters at 1, 3 and 5 dpf were compared to adult cells at 120 dpf **(Fig. 1F, S1D)**. EpGlia comprised 74.31% of total SC cells at 1 dpf **(Fig. 1F)**, reflecting the highly proliferative neuroepithelial cells that prevail during early development. In 3 and 5 dpf larvae, neurons predominated and comprised 60.30% and 59.07% of total SC cells, respectively. Compared to embryos and larvae, adult SC tissues at 120 dpf elicited a more balanced cell distribution between neurons, OPC/Oligos and EpGlia. Consistent with the developmental timeline, the proportions of EpGlia decreased from 74.31% at 1 dpf to 28.59% at 5 dpf. Conversely, neurons and immune cell proportions increased 5-fold between 1 and 5 dpf. Relative to 1 dpf, OPC/Oligos showed a more scattered UMAP distribution at 3 and 5 dpf, an observation that likely reflects the switch from neurogenesis to oligogenesis at 3 dpf **(Fig. 1G)** (Ravanelli and Appel, 2015; Scott et al., 2020).

Neurons, OPC/Oligos and immune cell profiles gradually changed from 1 to 5 dpf. But the relative proportions were still markedly different between 5 and 120 dpf **(Fig. 1F, S1D-G, Table S2)**. In line with incomplete establishment of spinal cell types at 5 dpf, we noted that adult cell clusters exhibited greater transcriptomic variability and heterogeneity relative to larval cells **(Fig. 1G, S1E)**. This was most evident for adult neurons, which displayed larger Euclidean distances that spanned the circumference of larval neurons on the UMAP **(Fig. 1G, S1E)**. To better assess changes in gene expression for each cell population across time points, we computed similarity scores between cell clusters using “ClusterFold Similarity” **(Fig. 1H)** (Gonzalez-Velasco et al., 2022). This statistical method was recently developed to quantitatively evaluate similarities across independent datasets by bypassing the need for data correction or integration. The similarities obtained using this method are therefore expected to convey similarities independent of the tissues, species or sequencing methods (Gonzalez-Velasco et al., 2022). In this analysis, the transcriptional profiles of larval samples at 1, 3 or 5 dpf were relatively similar **(Fig. 1H)**. In contrast, adult EpGlia, neurons, OPC/Oligos and immune cells showed markedly low similarity scores with their larval counterparts at 1, 3 or 5 dpf **(Fig. 1H)**. These results suggested that SC development continues beyond 5 dpf and that adult spinal cells are more diverse and differentiated than larval cells.

### Development, homeostasis and injury exhibit distinct cell compositions

To determine the extent to which regeneration recapitulates development, we integrated transcriptomic data from 3 dpf larvae (“Larval”), uninjured adult SCs (“Adult”) and 7 dpi adult SCs (“Adult-SCI”) **(Fig. 2A)**. We chose 3 dpf as larval SCI studies have traditionally employed 3-4 dpf larvae for regeneration studies (Alper and Dorsky, 2022). To ensure all conditions are equally represented in this analysis, we down-sampled the Adult and Adult-SCI datasets to match the number of Larval cells (11,367 cells per condition), while preserving the relative proportion of each annotated cell type within each sample. Unsupervised clustering of this integration yielded 28 clusters, classified as EpGlia, neurons, OPC/Oligos, immune or pericytes **(Fig. 2B-C, S2A-C)**. Expression of canonical markers confirmed cluster identities, including *sncb* in neurons, *gfap* in EpGlia and *mpz* in OPC/Oligos **(Fig. 2B)**. Except for EpGlia showing comparable proportions across conditions, Adult and Adult-SCI samples displayed different cell composition compared to larvae **(Fig. 2C, S2D)**. Supporting extended oligogenesis and immune cell development, the proportions of OPC/Oligos and immune cells increased 2.36- and 3.16-fold, respectively, between Larval and Adult samples **(Fig. 2C)**. On the other hand, neurons comprised 53.89% of Larval SCs but their proportion decreased to 32.77% in Adult SCs **(Fig. 2C)**. Consistent with acute immune activation after SCI, immune cell proportions further increased from 8.95% in Adults to 21.61% in Adult-SCI at 7 dpi **(Fig. 2C)** (Shaw et al., 2024). These findings indicated dynamic changes in the composition of coarse cell clusters between Larval and Adult SCs, suggesting neurons, OPC/Oligos and immune cells are not fully established at 3 dpf.

**Figure 2.**
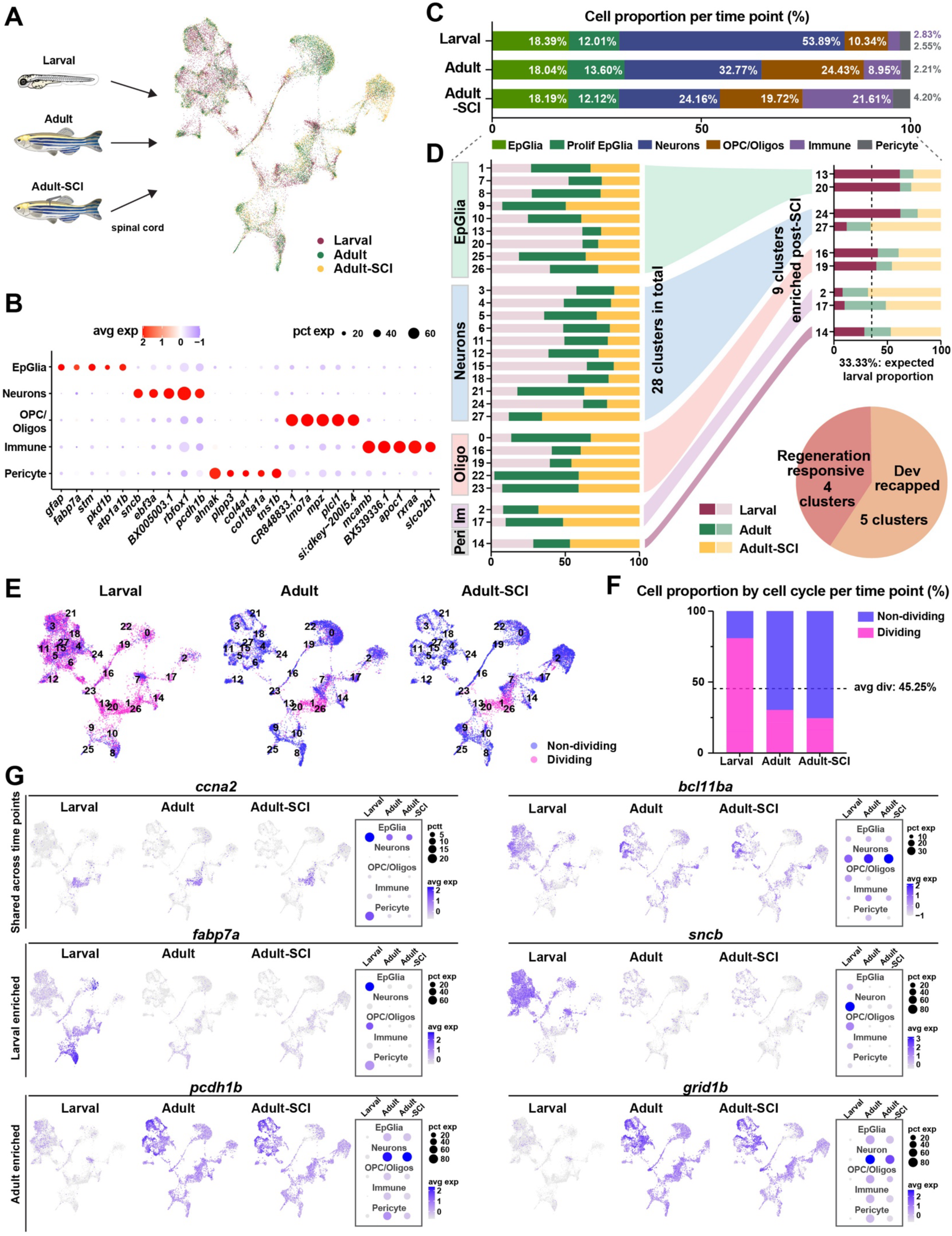
Assembly of a transcriptomic atlas for SC development, adult homeostasis and adult regeneration. **(A)** Integration of Larval (3 dpf), Adult (uninjured SCs at 120 dpf), and Adult-SCI (adult SCs at 7 dpi) samples. For larval analysis, only cells annotated as “neural”, “spinal cord”, “glial” or “immune” from 1, 3 and 5 dpf larvae were included. The adult datasets were generated from dissected SC tissues. **(B)** Dot plot shows the top 5 markers for each coarse annotation. Dot size and color represent percent expressing cells and average gene expression within each cell type, respectively. **(C)** Cell type composition of Larval, Adult and Adult-SCI samples. For each time point and coarse cluster, cell proportion was normalized to the total number of cells from that time point. **(D)** Distribution of major cell types across development and regeneration. The proportions of Larval, Adult or Adult SCI cells within each cluster is shown on the left. Clusters that increased by >25% in Adult-SCI compared to Adult samples were classified as injury-enriched and shown on the right. Among these injury-enriched clusters, the clusters that were comprised of >33.33% Larval cells were assigned as “development recapitulated” and represented in the pie chart. **(E)** Split UMAP representation of the integrated dataset, where dividing and non-dividing cells are highlighted. Cells in the G1 phase are annotated as non-dividing (blue), whereas samples in the G2/M and S phases are annotated as dividing (magenta). **(F)** Proportions of dividing and non-dividing cells within each time point of the integrated larval-adult dataset. **(G)** Feature plots and dot plot of markers that are i) shared across time points (*ccna2* and *bcl11ba*), ii) Larval-enriched (*fabp7a* and *sncb*), or iii) Adult-enriched (*pcdh1b* and *grid1b*).

To achieve a finer understanding of the cell dynamics that direct development versus regeneration, we surveyed the profiles of the 28 clusters from the integrated Larval, Adult and Adult-SCI dataset **(Fig. 2D, Fig. S2E)**. By comparing Adult versus Adult-SCI clusters, we first classified clusters that increased by >30% as “injury-enriched”. This analysis identified 9 injury-enriched clusters including EpGlia 13 and 20, neurons 24 and 27, OPC/Oligos 16 and 19, immune clusters 2 and 17, and pericyte cluster 14 **(Fig. 2D)**. Since total cell numbers were equivalent for each time point, we expected Larval cells would account for 33.33% of each cluster. Thus, clusters where Larval cells comprised >33.33% of cells within that cluster were considered “development recapitulated”. On the other hand, clusters with underrepresented Larval cells (<33.33%) were considered “regeneration responsive”. Notably, among the 9 injury-enriched clusters, over 50% of EpGlia 13, EpGlia 20 and neurons 24 were composed of Larval cells **(Fig. 2D)**. Conversely, some injury-enriched clusters such as neurons 27 were underrepresented in larvae. This analysis identified and distinguished between injury-responsive clusters that are likely to recapitulate development and injury-responsive clusters that specifically expand in adults after SCI.

### Development and regeneration activate distinct gene modules

To assess cell proliferation in larval and adult SCs, Seurat cell cycle scoring was used to categorize cells in the S, G1 or G2/M phases. Cells were annotated as dividing (S and G2/M) or non-dividing (G1) **(Fig. 2E)**. Overall, 81.38% of Larval cells and only 22.27% of Adult cells were dividing **(Fig. 2F)**. Following SCI, the proportion of proliferating cells was comparable between Adult and Adult-SCI samples. As Adult-SCI samples included 3 mm tissue blocks surrounding the lesion, it is possible that the scale of injury-induced proliferation was localized and diluted within the dissected tissue segments. An alternative possibility is that regenerative mechanisms maintain a controlled balance between cell proliferation and cell cycle exit for precise replenishment of lost cell types. Supporting the emergence of inherent proliferative differences between Larval and Adult samples, the Larval-predominant EpGlia clusters 20 and 26 were the most proliferative **(Fig. 2E, S2E-F)**. These results underscore further distinctions between the cellular landscapes that direct larval development and adult homeostasis.

We identified 3 gene expression motifs – genes shared across Larval and Adult samples (“Shared”), genes enriched in Larval samples (“Larval-enriched”), or genes enriched in Adult and Adult-SCI samples (“Adult-enriched”) **(Fig. 2G)**. The “Shared” gene expression motif included progenitor cell markers or drivers of neurodevelopment. For instance, *ccna2,* a marker of proliferating progenitors, was expressed in Larval and Adult SCs with higher baseline expression in larvae **(Fig. 2G)** (Cavone et al., 2021). *bcl11ba*, which encodes a transcription factor critical for neurodevelopment, was also expressed in Larval and Adult samples with preferential expression in adult neurons **(Fig. 2G)** (Lennon et al., 2017). On the other hand, “Larval-enriched” genes included *fabp7a* and *sncb* **(Fig. 2G)**. Consistent with the role of fatty acid binding protein in meeting the metabolic needs of multiple cell types, *fabp7a* was strongly expressed in Larval EpGlia, in addition to Larval OPC/Oligos and pericytes **(Fig. 2G)** (Pose-Mendez et al., 2024). *sncb*, which encodes the presynaptic phosphoprotein beta-Synuclein, was preferentially expressed in larval neurons **(Fig. 2G)** (Milanese et al., 2012). Finally, “Adult-enriched” genes included *pcdh1b,* which encodes a protocadherin that plays a key role in neuronal connectivity (Mincheva-Tasheva et al., 2024), as well as *grid1b,* which encodes a glutamate receptor subunit **(Fig. 2G)**. We propose genes enriched in Adult but not in Larval SCs are associated with more specialized neuromodulatory functions.

### Immune cell maturation is incomplete in larval zebrafish

Our coarse cluster analysis revealed drastic changes in the proportions and identities of immune cells and neurons between 3 dpf larvae and 120 dpf adults **(Fig. 2C, 3A, 3B)**. To delve into these differences, we subclustered immune cells and neurons from Larval, Adult and Adult-SCI datasets for further analysis **(Fig. 3C, 3D)**. The numbers of immune cells increased from Larval to Adult SCs and further increased in Adult-SCI samples at 7 dpi **(Fig. 3B)**. These observations are consistent with the extended timeline of immune cell maturation and with the emergence of an acute immune response after SCI (Willett et al., 1997; Davidson and Zon, 2004; Shaw et al., 2024). On the other hand, the numbers of neurons sharply declined from Larval to Adult SCs and decreased further after injury **(Fig. 3B)**. We postulated early neurogenesis is balanced by prolonged gliogenesis and immune cell maturation in juvenile zebrafish.

**Figure 3.**
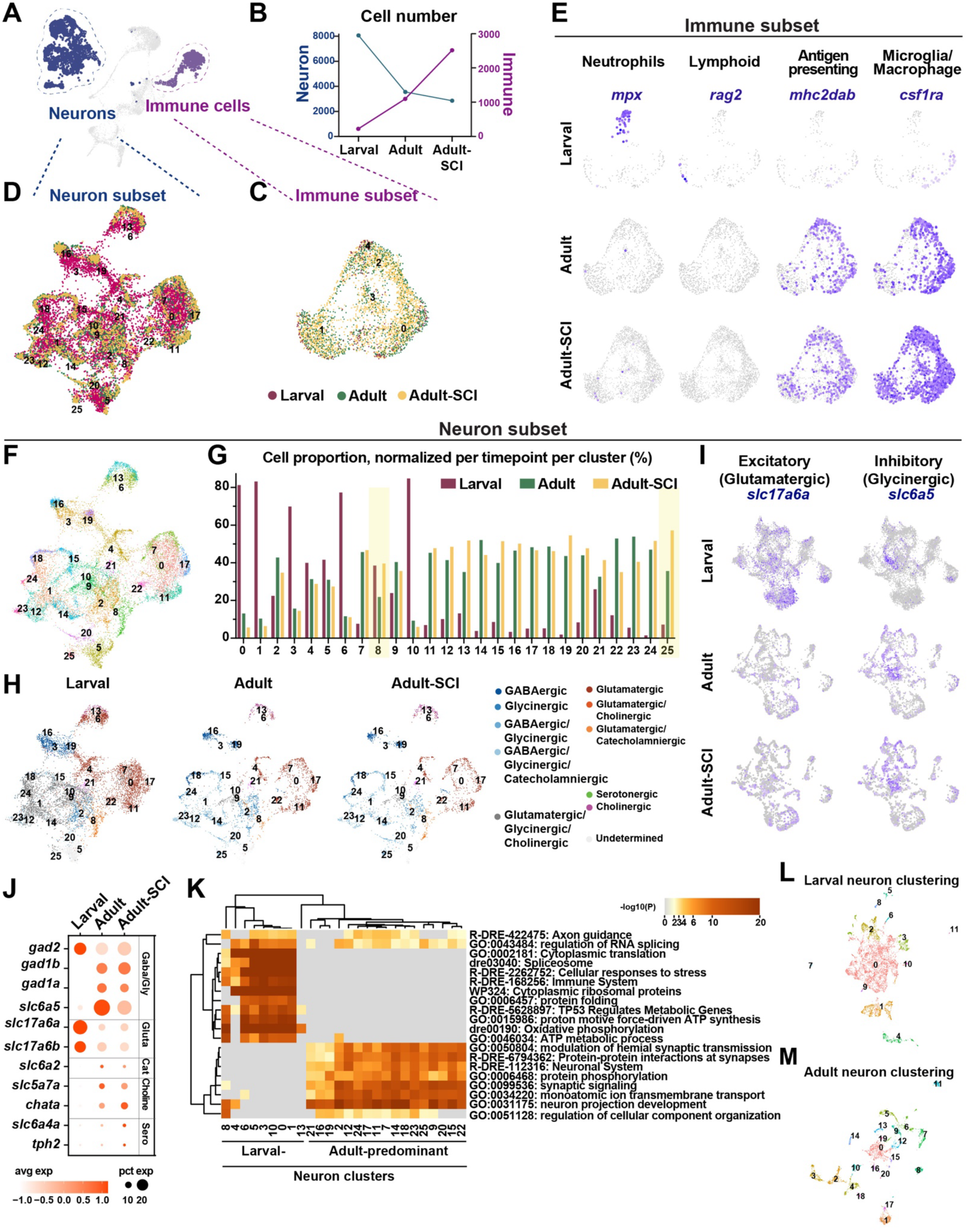
Adult SC regeneration deploys more mature immune cells and increased neuronal diversity compared to development. **(A)** UMAP representation of Larval (3 dpf), Adult (uninjured SCs at 120 dpf) and Adult-SCI (adult SCs at 7 dpi) samples. The highlighted immune cells (corresponding to clusters 2 and 15 from the integrated dataset in Fig. 2) and neurons (clusters 1, 3, 5, 10, 14, 16, 18, 19, 20, 21, 22, 23 and 25 from the integrated dataset in Fig. 2) are used for subcluster analyses. **(B)** Numbers of immune cells and neurons in Larval, Adult and Adul-SCI samples. **(C)** UMAP of immune cells after subsetting, normalization and integration. Cells are color-coded by time point. **(D)** UMAP of neurons after subsetting, normalization and integration. Cells are color-coded by time point. **(E)** Feature plots of immune cell markers within the immune subset. Markers shown from top to bottom are: *rag2* (lymphoid cell), *mpx* (neutrophils), *csf1ra* (microglia and macrophages) and *mhc2dab* (antigen-presenting cells). **(F)** UMAP of integrated neurons from Larval, Adult and Adult-SCI samples. Cells are color-coded by cluster number. **(G)** Neuronal cell proportions in Larval, Adult and Adult-SCI samples. For each cluster and time point, the numbers of cells per time point were normalized to the total number of cells in that cluster. Clusters that increased by >25% following SCI are highlighted in yellow. **(H)** Split UMAP of neuronal clusters, color-coded based on the neurotransmitter properties of each cluster. **(I)** Feature plots for the neuronal markers *slc6a5* (inhibitory glycinergic neurons) and *slc17a6a* (excitatory neurons) in Larval, Adult and Adult-SCI samples. **(J)** Dot plot of markers associated with GABAergic, Glycinergic, Glutamatergic, Catecholaminergic, Cholinergic and Serotonergic neurons in Larval, Adult and Adult-SCI samples. **(K)** Gene ontology for neuron subclusters in Larval, Adult and Adult-SCI samples. Represented on the heatmap are the most significantly enriched GO term for genes upregulated in each cluster. Columns represent cluster numbers. **(L-M)** Unsupervised Seurat clustering and UMAP representation were performed independently for Larval (L) and Adult (M) neurons from SC tissues respectively.

Despite their established roles in axon regrowth and neurogenesis after larval SCI (Tsarouchas et al., 2018; Cavone et al., 2021; de Sena-Tomas et al., 2024), immune cells were rare in developing larvae. To compensate for their reduced cell number and assuming that the blood-SC barrier is leaky at 3 dpf, we integrated all immune cells from whole larvae at 3 dpf (224 cells) with immune cells from dissected SC tissues of Adult (1,096 cells) and Adult-SCI (2,514 cells) samples **(Fig. 3C, S3A)**. Although UMAP visualization indicated good integration across samples **(Fig. 3C, S3B-E),** canonical marker gene expression revealed notable differences between Larval and Adult immune cells **(Fig. 3E)**. For instance, the lymphoid marker *rag2* and the neutrophil marker *mpx* were expressed in Larval but not Adult samples **(Fig. 3E)**. We postulate the presence of undifferentiated *rag2*^+^ lymphoid cells is likely due to the inclusion of hematopoietic stem cells outside the CNS or to an underdeveloped blood-SC barrier (Kvestak et al., 2024). On the other hand, *csf1ra^+^* microglia/macrophages and *mhc2dab^+^* antigen-presenting cells, which were almost absent in Larval tissues, accounted for most adult immune cells and expanded after injury **(Fig. 3E)**. Compared to larvae, adult immune cells displayed a more heterogenous cell composition, with fewer lymphoid cells, more microglia, macrophages and antigen-presenting cells. These findings suggested larvae and adults exhibit different immune profiles at baseline and are therefore likely to mount differential immune responses to SCI.

### Larval and adult neurons display distinct transcriptional profiles

To understand the transcriptional changes that support neuronal development and regeneration, we compared spinal neurons between Larval, Adult and Adult-SCI samples **(Fig. 3D)**. This neuronal clustering yielded 26 clusters **(Fig. 3F, S3F, S3G)**. Recapitulating our previous analysis from 1, 3, 5 and 120 dpf integration **(Fig. 1G)**, UMAP visualization showed greater transcriptional heterogeneity in Adult and Adult-SCI neurons compared to larvae **(Fig. S3F, S3G)**. Relative cell proportions validated that most neuronal clusters were predominately composed of either Larval neurons (e.g. clusters 0, 1, 3, 6 and 10) or Adult and Adult-SCI neurons (e.g. clusters 14, 16, 17, 18, 19, 23, 24, and 25) **(Fig. 3G, S3H)**. To survey the neurotransmitter properties of zebrafish neurons between development and regeneration, we annotated each cluster based on the expression of neurotransmitter-related genes **(Fig. 3H, S3I)**. This analysis included markers for glutamatergic (*slc17a6a*, *slc17a6b*), GABA/glycinergic (*slc6a5*, *gad1a*, *gad1b*, *gad2*), cholinergic (*chata*, *slc5a7a*), serotonergic (*slc6a4a*, *tph2*) and catecholaminergic neurons (*slc6a2*) **(Fig. 3I, S3J)**. We found Larval neurons exhibited a largely excitatory profile, enriched in glutamatergic markers and did not express other neurotransmitter-related genes **(Fig. 3H-J, S3J)**. Whereas most Larval neurons did not show a clear excitatory or inhibitory signature, Adult and Adult-SCI samples displayed more balanced and mature neurotransmitter profiles **(Fig. 3H-J)**. Gene Ontology of neuronal markers revealed a bimodal distribution, where terms associated with new cell synthesis and metabolism were expressed in Larval neurons while terms associated with neuronal activity were expressed in Adult neurons **(Fig. 3K)**. These findings indicated Larval and Adult neurons are intrinsically different in their heterogeneity, differentiation, neurotransmitter properties and gene expression profiles.

To confirm that Larval neurons at 3 dpf are less diverse and less differentiated than Adult neurons at 120 dpf, we performed an independent analysis that did not require any data integration. To this end, we clustered Larval and Adult neurons separately **(Fig. 3L, 3M)**. Clustering resolutions were selected based on the best balance between cluster granularity and the stability of top marker gene expression **(Fig. S3K, S3L)** (Zappia and Oshlack, 2018). Unsupervised clustering yielded 12 Larval and 21 Adult neuronal clusters. Moreover, Adult neurons displayed visibly distinct clusters, suggesting a more differentiated, mature neuronal profile compared to Larval neurons **(Fig. 3L, 3M)**. Thus, regardless of sequencing or integration methods, Adult SC neurons exhibit greater heterogeneity than Larval neurons, indicating prolonged neuronal maturation during larval and juvenile growth.

### The molecular profiles of *sox2^+^* cells between development and adult homeostasis

*sox2* expression marks SC progenitors during vertebrate development and regeneration (Fei et al., 2014; Ogai et al., 2014; Hui et al., 2015; Donato and Vickaryous, 2022). To investigate the molecular identities of SC progenitors, we subclustered *sox2^+^* cells from Larval, Adult and Adult-SCI samples. *sox2^+^* cells comprised 15.06% of Larval, 2.45% of Adult and 6.91% of Adult-SCI samples **(Fig. 4A, S4A)**. To analyze *sox2^+^* progenitors independently of their sample size differences, we down-sampled *sox2^+^* cells from Larval and Adult-SCI cells to match the number of Adult progenitors (808 *sox2*^+^ cells per sample). This analysis yielded 13 clusters with distinct molecular identities reflecting their heterogeneity and/or lineage biases **(Fig. 4B)**. We have previously optimized cell type identification by custom-building a database of CNS markers from over 12 datasets and vertebrate species including zebrafish, mice and humans (Saraswathy et al., 2024). To determine the identities and lineage biases of *sox2^+^*cells, we cross-referenced differentially expressed (DE) markers for each *sox2^+^* cluster with our marker database. Scoring matrix heatmap from this analysis identified biases toward EpGlia, neurons or OPC/Oligos **(Fig. S4B-E)**. Reflecting active neurogenesis during development, the majority of *sox2*^+^ cells in the Larval sample were neuron-biased, ostensibly including progenitors that are primed toward neuronal differentiation as well as newborn neurons that maintained remnant *sox2* expression **(Fig. 4C)**. Neuron-biased progenitors decreased from 59.16% in Larval to 24.38% in Adult and 19.43% in Adult-SCI samples **(Fig. 4C)**. On the other hand, the proportions of OPC/Oligo-biased *sox2^+^* cells increased from 4.83% in Larval to 26.24% in Adult and 42.57% in Adult-SCI tissues **(Fig. 4C)**. These numbers suggested that OPCs expand throughout larval and juvenile growth and that some OPC/Oligos upregulate *sox2* expression following SCI. These findings also suggested distinct neurogenic progenitors may underlie SC development and adult regeneration.

**Figure 4.**
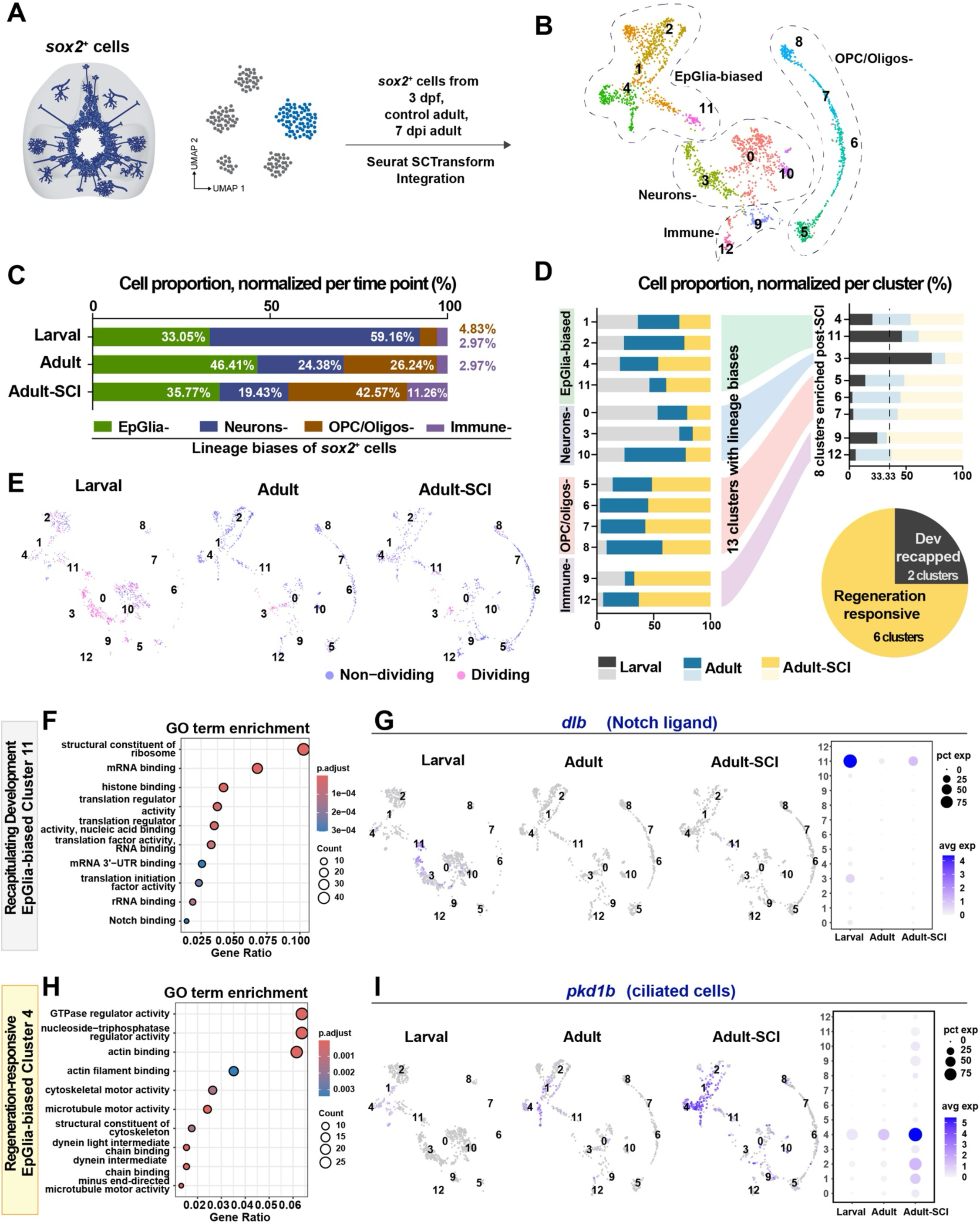
Subclustering *sox2*^+^ progenitors during SC development and adult regeneration identifies regeneration-specific clusters. **(A)** Schematic representation of *sox2*^+^ cells (blue) in SC cross sections following SCI. *sox2*^+^ cells were from Larval, Adult and Adult-SCI samples were integrated for subsequent analysis. **(B)** Combined UMAP of *sox2*^+^ cells from Larval, Adult, and Adult-SCI SCs, annotated based on their lineage biases. **(C)** Representation of lineage biases in *sox2*^+^ cells from Larval, Adult and Adult-SCI samples. For each lineage bias and time point, cell proportions were normalized to the total number of cells for that time point. **(D)** Distribution of major lineage biases during development and regeneration. The proportions of Larval, Adult or Adult SCI *sox2*^+^ cells within each cluster is shown on the left. Clusters that increased by >25% in Adult-SCI compared to Adult samples were classified as injury-enriched and shown on the right. Among these injury-enriched clusters, the 2 clusters that were comprised of >33.33% Larval cells were assigned as “development recapitulated” and represented in the pie chart. The remaining 6 clusters were considered “regeneration responsive”. **(E)** Split UMAP of *sox2^+^* cells shows dividing and non-dividing cells. Cells in the G1 phase are annotated as non-dividing (blue), and samples in the G2/M and S phases are annotated as dividing (magenta). **(F)** Dot plot representing the top enriched GO terms for cluster 11, which recapitulates development. **(G)** Feature plot of *dlb*, a top marker of cluster 11. **(H)** Dot plot representing the top enriched GO terms for cluster 4, which is regeneration-responsive. **(I)** Feature plot of *pkd1b*, a top marker of cluster 3.

### Distinct pools of *sox2^+^* cells are activated during development and regeneration

Out of 13 *sox2*^+^ cell clusters, 8 clusters expanded >33% in Adult-SCI compared to Adults and were categorized as injury-enriched **(Fig. 4D)**. Injury-enriched clusters included EpGlia-biased clusters 4 and 11, neuron-biased cluster 3 and OPC/Oligos-biased clusters 5, 6 and 7 **(Fig. 4D)**. To determine if these injury-enriched signatures recapitulate progenitor cell dynamics during development, we examined the proportions of Larval cells within each cluster. Since total cell numbers were equivalent for each time point, clusters where Larval cells comprised >33.33% of total cells within that cluster were considered “development recapitulated”, whereas clusters with underrepresented Larval cells (<33.33%) were considered “regeneration responsive”. This analysis showed only 2 of the 8 (25%) injury-enriched clusters showed similar transcriptional profiles to larval progenitors (clusters 3 and 11) **(Fig. 4D)**. Conversely, Larval cells were underrepresented in 6 of the 8 (75%) injury-enriched clusters, suggesting the molecular signatures associated with SC regeneration are largely injury-responsive and not a faithful recapitulation of development.

We observed varying cell cycle states across clusters **(Fig. 4E)**. The proportion of dividing *sox2*^+^ cells was highest during development, accounting for 65.22% of Larval cells **(Fig. S4F)**. On the other hand, only 14.24% and 15.47% of Adult and Adult-SCI cells were classified as dividing **(Fig. S4F)**. Neuron-biased *sox2*^+^ cluster 3 displayed a notably high rate of cell division (93.78%) and expressed *pcna* as a top marker **(Fig. S4G)**. Among the 4 EpGlia-biased clusters, only cluster 11 exhibited increased proliferation. Intriguingly, despite different lineage biases, clusters 11 and 3 showed relatively adjacent transcriptome signatures on the UMAP, forming a “bridge” between EpGlia- and neuron-biased progenitors **(Fig. 4B)**. Clusters 11 and 3 were also the only injury-enriched clusters that recapitulated development **(Fig. 4D)**. We postulate that clusters 3 and 11 comprise transient populations of *sox2^+^*progenitors that redeploy developmental pathways to direct neurogenesis following SCI.

To understand the molecular differences between injury-enriched clusters that recapitulate development and ones that are adult-specific, we delved into the transcriptional signatures of EpGlia-biased clusters 11 and 4 **(Fig. 4F-I)**. Cluster 11 is an injury-enriched cluster that recapitulates development **(Fig. 4D)**. GO analysis showed genes associated with Notch signaling including *dlb, dla* and *her4.2* were enriched in cluster 11 in addition to terms associated with cellular component synthesis **(Fig. 4F, S4H, Table S3)**. Consistent with Notch reactivation during regeneration, *dlb* was a top marker of cluster 11 and displayed higher expression in Larval samples compared to either Adults or Adult-SCI **(Fig. 4G)**. On the other hand, cluster 4, which was underrepresented in Larval tissues **(Fig. 4D),** expressed genes implicated in cytoskeletal dynamics, microtubule, or actin motor activities **(Fig. 4H, S4H, Table S3)**. Consistent with a ciliated morphology, the top marker for cluster 4 was the ependymal marker *pkd1b* **(Fig. 4I, S4H, Table S3)**. These studies highlighted the heterogeneity of *sox2^+^* cells during development and regeneration, and identified a cluster of EpGlia-biased progenitors that are likely to convey pro-regenerative potential to adult zebrafish after SCI.

### Maturation and positional identity are transcriptionally encoded in larval SC progenitors

SC progenitors are patterned into spatially and temporally defined domains during development (Sagner and Briscoe, 2019). Spatially, opposing gradients of dorsal BMP and ventral SHH signaling establish coarse dorso-ventral (D-V) progenitor domains (Wilson and Maden, 2005; Kicheva and Briscoe, 2023). Temporally, early morphogen-regulated transcription factors encode gene regulatory networks that further diversify D-V cell identities at later developmental stages (Osseward et al., 2021). Using the high-resolution transcriptome of larval chronological ages from DanioCell, we characterized the maturation and spatial identities of SC progenitors between 14 and 120 hpf. To capture spinal EpGlia from DanioCell, we first verified *hox* gene expression across “tissue.clust” annotations. *hoxb8a,* which is expressed in developing SC tissues, was expressed in most “glial” and “SC” clusters, in addition to 1 “neural” cluster (Holstege et al., 2008). By cross-referencing the identities of *hoxb8a^+^* clusters to the DanioCell identity annotations, we filtered out non-SC neural cells. These steps generated a temporally continuous subset of larval EpGlia from 14 to 120 hpf.

To determine whether “Maturation” of progenitor cells could be extrapolated from this transcriptional dataset, we subclustered *hoxb8a^+^* EpGlia between 14 and 120 hpf **(Fig. S5A).** We then annotated each cell on the UMAP based on the chronological age of the sample it was collected from **(Fig. 5A)**. Noting that progenitor cells at 14 hpf and 120 hpf were the most distant on the UMAP with an observable gradient of intermediate time points, we concluded that the chronological age of progenitor cells is indeed encoded in their transcriptional signatures **(Fig. 5A)**. To confirm whether “D-V” identities are also encoded in the transcriptome of progenitor cells, we next compiled a list of D-V markers that are known to mark neural tube development (Wilson and Maden, 2005). Markers including *shha, ptch2, nkx2.2b, olig2, nkx6.1, pax3a, msx1a, msx1b, pax6a, pax7a, irx3a, lbx2, nkx6.2, gsx1* and *dbx1b* were used **(Table S4)**. Remarkably, clustering larval EpGlia using these D-V markers as variable features recapitulated their anatomical location along the D-V axis of zebrafish larvae **(Fig. 5B)**. This analysis established a reference atlas of progenitor cell maturation and D-V identities during larval SC development.

**Figure 5.**
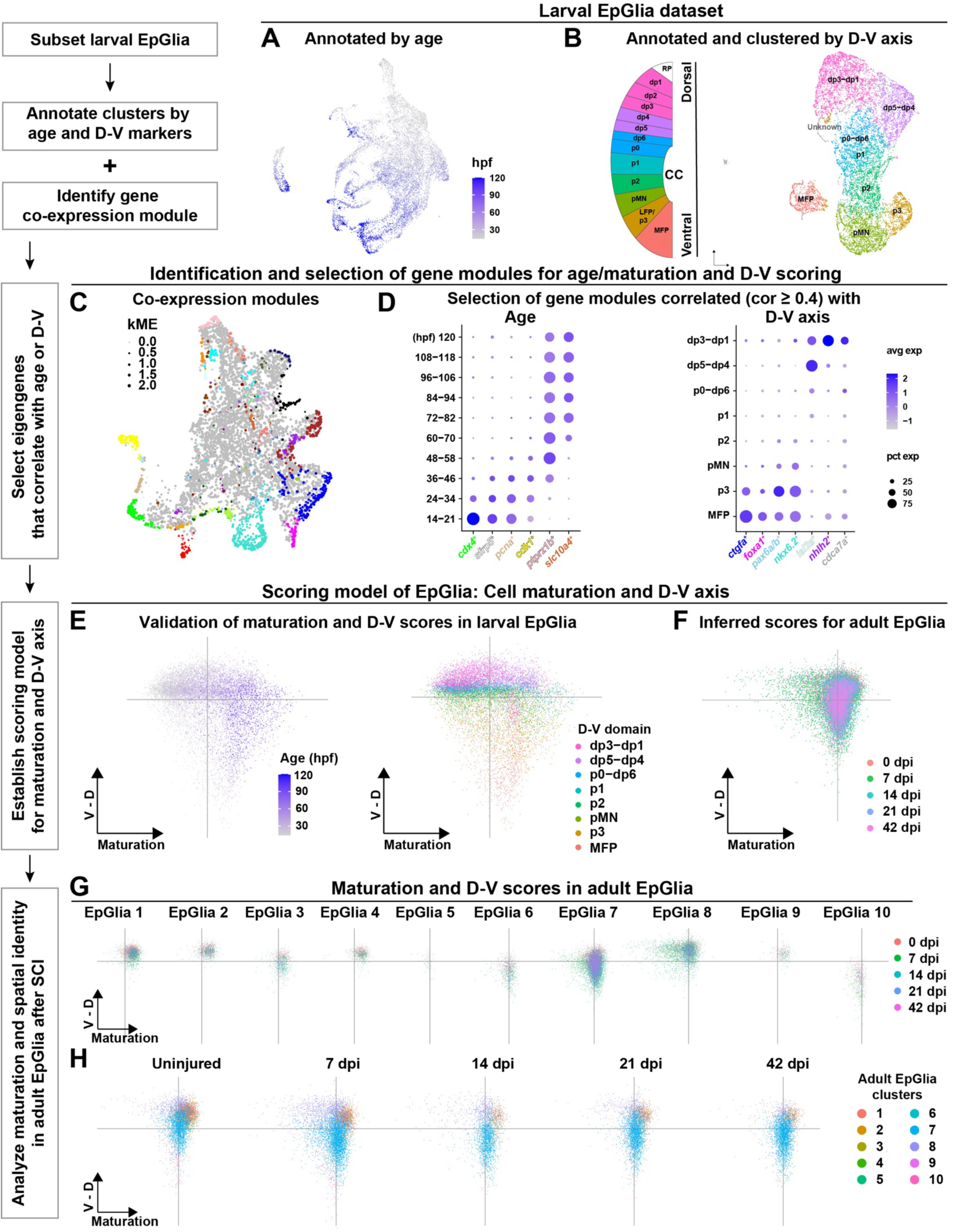
The dorso-ventral progenitor identities that guide SC development are not maintained in adult regeneration. **(A)** Unsupervised Seurat clustering and UMAP representation of larval EpGlia between 14 to 120 hpf. Cells are colored based on the chronological stage of the samples they were collected from. **(B)** Larval EpGlia were clustered using a series of progenitor markers along the D-V axis. The resulting clusters showed marked parallels with their D-V anatomical location in the developing neural tube. CC: central canal. **(C)** UMAP of gene modules identified from larval EpGlia. Each node represents a single gene and point size was scaled by kME. Each color represents one co-expression module. Grey nodes are unclassified genes. **(D)** Dot plot of larval EpGlia gene co-expression modules that correlate with chronological age (left) or with D-V position (right). These eigengenes were used to establish maturation and D-V scores. **(E)** Validation of our maturation and D-V scoring system in larval EpGlia. PC plots of larval EpGlia between 14 to 120 hpf are shown. Cells were color-coded based on the larval time point at which they were collected (left) or their D-V position (right). **(F)** Inferred maturation and D-V scores for adult EpGlia. PC plot of adult EpGlia from 0, 7, 14, 21 and 42 dpi is shown. Scalar multiplication was performed using the D-V and maturation scoring matrices established for larval EpGlia. Cells were color-coded based on the time point at which they were collected. **(G)** PC plots of Adult EpGlia clusters based on our larval maturation and D-V scoring system. PC plots are split per EpGlia clusters and cells are color-coded based on the time point at which they were collected. **(H)** PC plots of Adult EpGlia clusters based on our larval maturation and D-V scoring system. PC plots are split per time point and cells are color-coded based on their EpGlia cluster classification.

### Scoring “Maturation” and “D-V” identities in larval SC progenitors

We identified major gene expression modules (i.e. eigengenes) of larval SC progenitors using the “hdWGCNA” package. With a minimum of 5 genes set for module classification, a total of 46 modules were identified **(Fig. 5C)**. To determine gene modules that correlate with either developmental age or D-V axis while eliminating compounding correlation between these two factors, we computed correlation factors for each module with either chronological age or D-V identity **(Fig. S5B)**. We then selected gene modules with correlation larger than the absolute value of 0.4 and filtered out overlapping modules. This analysis revealed 6 age-related and 7 D-V-related eigengenes **(Fig. 5D)**. We validated that larval EpGlia collected at increasing chronological age displayed gradually higher maturation scores that stabilized around 72 hpf **(Fig. S5C)**. We also confirmed the D-V scores of larval EpGlia matched their D-V axis location **(Fig. S5D)**. Using maturation and D-V scores as principal components, we then performed PCA analysis on larval EpGlia **(Fig. 5E)**. By pseudo-coloring EpGlia based on either the chronological ages of the embryos they were collected from or their D-V location, we validated the distribution of early-to-late and ventral-to-dorsal EpGlia along the X and Y axes, respectively **(Fig. 5E)**. This analysis established a scoring model to assess maturation and D-V identities in SC progenitors.

### Adult SC progenitors do not maintain the dorso-ventral identities of larval progenitors

To investigate whether adult SC progenitors recapitulate the temporal and spatial signatures of larval progenitors, we performed scalar multiplication on each adult EpGlia sample based on the reference PCA values of larval EpGlia **(Fig. 5F)**. To maximize the capture of transitional states during SC regeneration, adult EpGlia from multiple SCI time points (7, 14, 21 and 42 dpi) in addition to uninjured SC tissues were included (Saraswathy et al., 2024). This analysis clustered adult SC progenitors into 10 EpGlia clusters, revealing several differences in maturation and D-V axis distribution between larvae and adults **(Fig. 5F-H)**. Compared to their uninjured counterparts, adult EpGlia exhibited a modest and transient shift to less mature states and more ventral identities at 7 dpi **(Fig. 5G, 5H)**. Except for clusters 6, 7 and 10 showing a more ventral distribution compared to other clusters, adult EpGlia clusters did not show a clear stratification along the D-V axis **(Fig. 5G)**. EpGlia clusters 1-4 scored high in cell maturation, while EpGlia clusters 7 and 8 exhibited low maturation scores and high variance post-injury **(Fig. 5G, 5H)**. These changes suggested clusters 7 and 8 transition into an immature state early after SCI and become more mature at later stages of regeneration. Nonetheless, compared to larval progenitors that extended into precisely mapped spatio-temporal domains **(Fig. 5E)**, adult EpGlia exhibited a more compressed distribution on the maturation and D-V axis **(Fig. 5G)**. These findings suggested the transcriptional principles that govern larval progenitor identities may not be necessarily conserved in adult progenitors.

We validated the reference PCA graph of Larval EpGlia by surveying canonical marker gene expression **(Fig. 6A)**. For instance, ventrally expressed genes such as *shha* and *foxa1*, dorsally expressed genes such as *pax3a* and *irx3a*, as well as early progenitor markers such as *msx1a* and *msx1b* recapitulated their temporal features and anatomical locations on the PCA graph **(Fig. 6A)**. By visualizing the same canonical markers in adult progenitors, the distribution and boundaries of the D-V domains were not clearly stratified **(Fig. 6B)**. Markers such as *prdm12b* and *msx1a*, which are spatially restricted to p1 and dorsal domains in larvae, showed sparse expression in adult EpGlia **(Fig. 6A, 6B)** (Liu et al., 2004; Zannino et al., 2014). Intriguingly, while the expression of the *dp1-3* marker *pax3a* was dorsally restricted in the larval dataset, *pax3a* expression was widely distributed along the D-V axis in adult EpGlia **(Fig. 6A, 6B)**. Finally, expression of the medial progenitor marker *dbx1b* was restricted to a band-like stratification on the D-V axis but spanned cells of all maturity scores in Larval EpGlia **(Fig. 6A)** (Briona and Dorsky, 2014). *dbx1b* expression in Adult EpGlia exhibited a wider D-V distribution but was more restricted on the maturation axis **(Fig. 6B)**. This analysis showed adult spinal progenitors loosely captured but did not faithfully recapitulate the dorso-ventral identities of larval EpGlia, suggesting spatial precision may be less required during regeneration than development.

**Figure 6.**
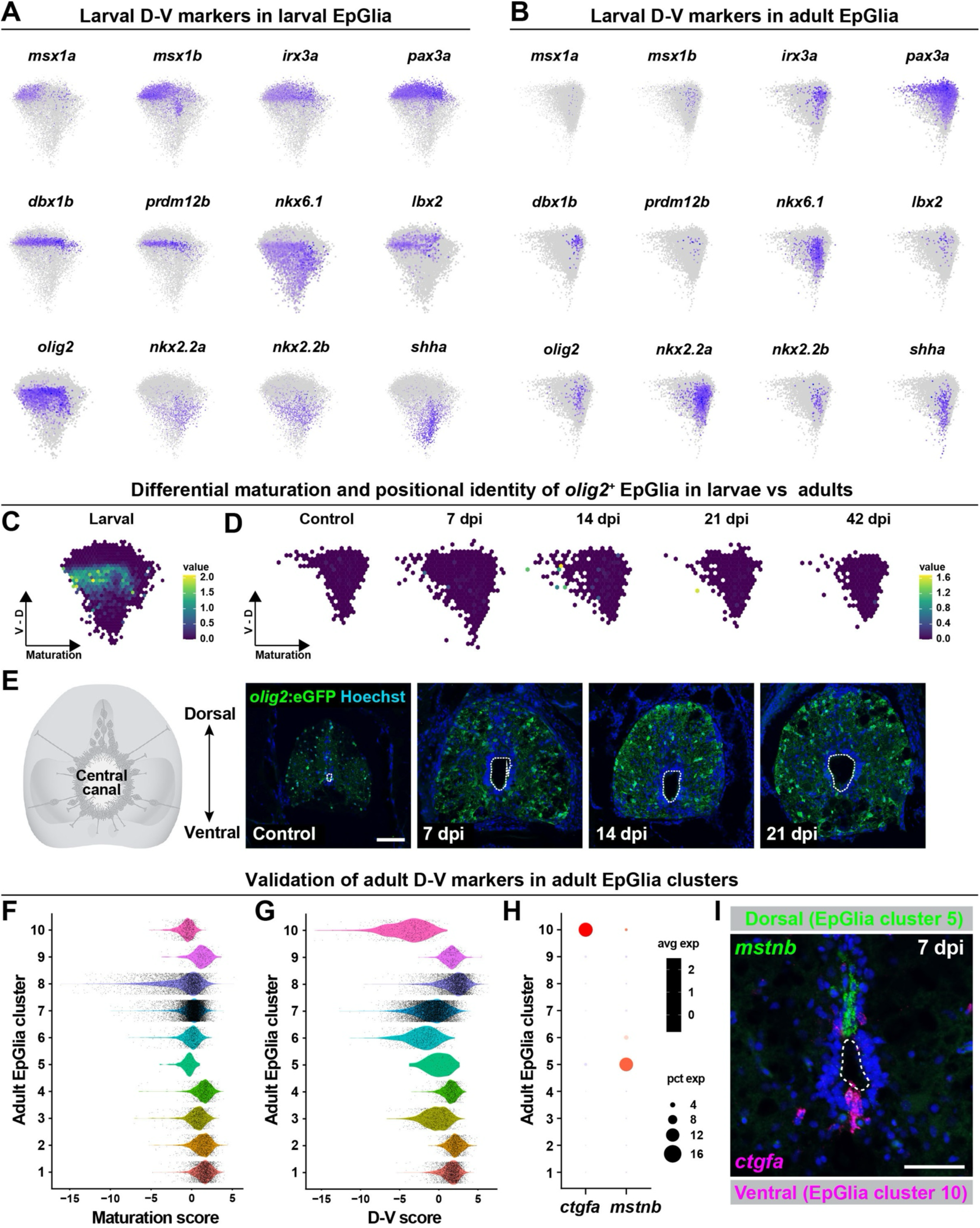
Validation of larval and adult D-V markers in adult in adult SC progenitors. **(A)** Validation of larval D-V markers in larval EpGlia. Feature plots show select larval D-V identity genes. PC plots of larval EpGlia between 14 and 120 hpf are shown. **(B)** Validation of larval D-V markers in adult EpGlia. Feature plots show select larval D-V identity genes. PC plots of adult EpGlia from 0, 7, 14, 21 and 42 dpi are shown. **(C, D)** Feature plots show *olig2* expression in larval (C) and adult (D) EpGlia. **(E)** HCR *in situ* hybridization validates that the dorso-ventral spatial identities of larval *olig2*-expressing cells are not maintained in adult homeostasis or regeneration. HCR was performed on SC sections from *olig2:GFP* transgenic line at 0, 7, 14 and 21 dpi. Dotted lines delineate the central canal. **(F, G)** Violin plots show our calculated maturation scores (F) and D-V scores (G) for each adult EpGlia cluster. **(H, I)** *in vivo* validation of D-V clusters 5 and 10 in adult SC tissues. Dot plot (H) and dual HCR *in situ* hybridization (I) show *mstnb* and *ctgfa* expression in dorsal EpGlia cluster 5 and ventral EpGlia cluster 10, respectively. HCR was performed on wild-type SC sections at 7 dpi. Dotted lines delineate the central canal. Scale bars: 50 μm.

### *in vivo* validation of larval and adult progenitor markers in adult SC progenitors

Progenitor cells expressing *olig2* direct sequential generation of motor neurons and oligodendrocytes during embryonic and larval development (Park et al., 2007; Ravanelli and Appel, 2015). Our maturation and D-V scoring showed *olig2*^+^ cells possess varied maturation and D-V scores in larval and adult EpGlia **(Fig. 6C, 6D)**. Larval *olig2* expression was restricted to less mature progenitors of the pMN domain that also express *nkx6.1* **(Fig. 6A, 6C)**. Conversely, *olig2* expression in adult EpGlia showed higher maturation scores and was predicted to span a wider range of D-V localization **(Fig. 6B, 6D)**. To validate these *in silico* predictions, we assessed GFP expression in adult zebrafish harboring an *olig2:eGFP* transgene at 7, 14 and 21 dpi in addition to uninjured adults. in adult SC tissues. In SC cross sections, *olig2:eGFP* expressing cells were uniformly distributed around the parenchyma in adult homeostasis and after SCI **(Fig. 6E)**. These findings suggested that canonical D-V markers of larval EpGlia do not maintain restricted spatial identities in adult SC tissues.

To examine whether adult EpGlia possess molecular identities specific to adult D-V regions, we examined adult EpGlia clusters along the maturation and D-V axes **(Fig. 6F, 6G)**. We noted adult EpGlia cluster 10 expressed *connective tissue growth factor a (ctgfa)* and scored the lowest on D-V axis **(Fig. 6G, 6H)**. Intriguingly, *ctgfa* expression is a hallmark of ventral progenitors after adult SCI, and was shown to be required for glial bridging in adult zebrafish but dispensable in larvae **(Fig. 6H)** (Mokalled et al., 2016; Wehner et al., 2017; Zhou et al., 2023; Walker et al., 2025). Conversely, adult EpGlia cluster 5 exhibited a higher dorsal score and expressed *myostatin b (mstnb)* **(Fig. 6G, 6H)**. *mstnb* expression marks dorsal progenitors after adult SCI and was shown to modulate the rates of self-renewal and neurogenesis in dorsal SCs (Saraswathy et al., 2022). Validating our *in silico* predictions, dual HCR *in situ* hybridization showed *ctgfa* and *mstnb* are expressed at 7 dpi in distinct niches of adult SC progenitors in ventral and dorsal SC tissues, respectively **(Fig. 6I)**. These findings suggested that canonical D-V markers of larval EpGlia do not maintain restricted spatial identities in adult SC tissues and that adult EpGlia express molecular identity genes that do not necessarily mirror larval D-V identities.

## DISCUSSION

This study assembles a single-cell atlas of SC development and regeneration in zebrafish, providing insights into the cell compositions and molecular machinery that direct development versus regeneration. By integrating cell identity features including maturation and dorso-ventral identity, we found the spatio-temporal progenitor identities that are characteristic of SC development are not maintained in regenerating adults.

The advent of single-cell transcriptomics has significantly advanced the resolution of our molecular studies, yielding new insights into SC development and regeneration (Farrell et al., 2018; Cavone et al., 2021; Scott et al., 2021; Sur et al., 2023; Lange et al., 2024; Saraswathy et al., 2024). Realizing our single-cell (larval) and single-nuclear (adult) integration may be cofounded by batch differences, we implemented multiple quality checks to rule out technical artifacts (Sur et al., 2023; Saraswathy et al., 2024). First, our results are based on multiple, independent integrations using various sample combinations, controlled cell numbers and rigorous integration parameters. As all iterations yielded good integration among biological samples and reproducible observations, we expect technical or batch effects to be limited. Importantly, our results were validated using integration-independent tools. For instance, “ClusterFold Similarity”, which bypasses the need for data correction or integration, revealed pronounced differences between larval and adult cells (Gonzalez-Velasco et al., 2022). Moreover, independent clustering of larval and adult neurons yielded more adult clusters. These results support a model in which SC development continues beyond 5 dpf, whereby adult neurons are more diverse and specified than larval neurons. In alignment with recent integrations of scRNA-seq and snRNA-seq datasets (Quatredeniers et al., 2023), we anticipate that integrative comparative transcriptomics will continue to yield valuable insights into various developmental and regenerative systems.

Regenerative vertebrates harbor potent SC progenitors that underlie their regenerative capacity. Our study employs 2 complementary methods to unravel the molecular identities and developmental origins of adult SC progenitors. We first asked whether adult progenitors are residual embryonic radial glia that persist in adult SC tissues, or a hybrid population of adult ependymal cells that retain radial glia features. Integrated analysis of *sox2*^+^ cells from 3 or 5 dpf larvae and 120 dpf adults showed distinct molecular identities with only 25% of injury-enriched adult clusters recapitulating development. In fact, even within the clusters that recapitulate development (clusters 3 and 14 in Fig. 4C-D), larval and adult cells showed slightly different UMAP distribution. Validating these findings, independent analysis of chronological age and D-V identities yielded further differences between larval and adult EpGlia. Whereas the D-V identities characteristic of SC development were altered in adult EpGlia clusters, the D-V clusters that were identified in this analysis were preferential to adult SC regeneration. Notably, ventral and dorsal EpGlia clusters based on D-V scoring expressed *ctgfa* and *mstnb*, which were reported to direct glial and neuronal repair during adult SC regeneration (Mokalled et al., 2016; Klatt Shaw et al., 2021; Saraswathy et al., 2022; Zhou et al., 2023). Consistent with the emergence of adult-specific regenerative mechanisms, *ctgfa*-driven glial bridging is required for glial bridging in adult zebrafish and 7 dpf larvae, but not in younger larvae (Briona et al., 2015; Mokalled et al., 2016; Wehner et al., 2017; Klatt Shaw et al., 2021; Zhou et al., 2023; Walker et al., 2025). We propose that in depth molecular evaluation of various progenitor clusters is needed to determine whether a subset of adult progenitors are true ependymal-radial glial hybrid cells.

Our study reports SC tissues harbor distinct cell identities between 3 or 5 dpf larvae and 120 adults. These findings are inconsistent with previous reports that larval SC progenitors become fate-restricted at 3 dpf, and that zebrafish larvae establish all CNS cell types at that time point (Parichy et al., 2009; Alper and Dorsky, 2022). These results are also consistent with recent single-cell integrations between age-matched developing and regenerating SC tissues in Xenopus (Swearer et al., 2025). Crucial for SCI research, both larval and adult zebrafish are increasingly (and sometimes interchangeably) used to uncover fundamental neural repair mechanisms. Thus, as baseline cell composition and molecular identities are different between larval and adult homeostatic SCs, the quality and/or magnitude of some injury responses may differ between these models. Moreover, unlike the continuous progression of embryonic development, adult regeneration is triggered by the occurrence of an abrupt injury that activates dormant injury responsive programs. As numerous developmental pathways show baseline activity in 3 dpf larvae, it is possible that the burdens to activate regenerative programs are less pronounced in larval SCI. Additional direct comparative studies between larval and adult zebrafish SCI are needed to determine whether larval SCI is a closer recapitulation of larval development or adult SCI. Despite their differences, both larval and adult zebrafish are valuable models to understand fundamental principles of innate SC regeneration, and we encourage researchers to harness the unique advantages of each model to address their scientific inquiry.

## ACKNOWLEDGMENTS

We thank V. Cavalli, A. Johnson and P. Williams for discussion. This research was supported by grants from the NIH (R01 NS113915 to MHM) and the Irving Boime Graduate Fellowship from the Department of Developmental Biology at Washington University School of Medicine (to YX).

## METHODS

### sc/snRNA-seq data acquisition

Multiple integrations were performed using previously published sc/snRNA-seq data. scRNA-seq data from whole zebrafish embryos/larvae between 14 and 120 hpf were obtained from DanioCell (Sur et al., 2023). In this whole embryo dataset, SC cells were broadly identified based on the expression of CNS markers. To distinguish between brain and SC cells, anterior-posterior markers during neurodevelopment (*otx2a, otx2b, emx1, hoxa1a, hoxb1a, hoxb2a, hoxa2b, hoxc6a, hoxb6b, hoxb8a, hoxa9a, hoxc9a, and hoxa11b*) were visualized. Samples from “neural”, “SC” and “glial” annotations on the “tissue.figure” identities were included to ensure a comprehensive comparison of SC tissues from development to adulthood. All cells annotated as “immune” from the larval dataset were included. snRNA-seq data from adult zebrafish SCs were previously published (Saraswathy et al., 2024). The adult dataset was collected from SC segments spanning 3 mm around the lesion site.

### Data integration and Seurat analysis

Datasets were integrated and analyzed using Seurat (v4) package with R (v4.3.1). Each dataset was independently normalized and scaled using the “SCTransform” function. Standard Seurat integration workflow was used to identify shared sources of variation across experiments as well as mutual nearest neighbors. Integration features were selected based on the top 2000 variable features using “SelectIntegrationFeatures” function (nfeatures = 2000), which was used as input for the “anchor.features” argument of the “FindIntegrationAnchors” function. PCA analysis was performed on the variable features. The top 50 principal components were selected based on the elbow plot heuristic, which measures the contribution of variation in each component. These 50 principal components were used in “FindNeighbors” and “FindClusters” functions to perform graph-based clustering on a shared nearest neighbor graph. Louvain algorithm was used for modularity optimization in cell clustering using “FindClusters” function. The resolution parameter that determines the granularity of clustering was selected by visually inspecting clusters with resolutions ranging between 0.1 and 2.0, as well as clustree graphs (Zappia and Oshlack, 2018). Uniform Manifold Approximation and Reduction (UMAP) representation was used for non-linear dimensional reduction of the first 50 principal components and data was visualized using “RunUMAP” function. Data was graphed using different plot packages and functions, such as “plot1cell”, “DimPlot”, “VlnPlot”, “FeaturePlot” and “Dotplot” to view cluster identity and marker gene expression. Cell numbers were extracted using the “table” function. Cell proportions were then calculated and graphed using GraphPad Prism. Raw cell numbers and normalized cell proportions were reported.

### Identification of differentially expressed (DE) markers

Differential gene expression for individual cell clusters was determined using Wilcoxon rank sum test in the “FindAllMarkers” function. Marker genes detected in at least 10% of the grouped cells and with a logFC threshold of 0.25 were selected. Only positive markers were reported.

### Cell type and lineage bias identification

A previously compiled “CNS marker” database was first used to identify cell types and lineage-biases based on top DE markers (Saraswathy et al., 2024). For each cluster, the top differentially expressed (DE) marker genes of that cluster were cross-referenced with the “CNS marker” database. A scoring matrix was generated and normalized to the total number of markers available in the database under each cell type category. The resulting values in the scoring matrix were scaled to 0-100 and plotted as a heatmap using GraphPad Prism. Each cluster was given an identity based on the maximum score obtained in the heatmap. To confirm the assigned identity, expression of canonical cell markers was evaluated. To further validate cell type and lineage bias annotations, the cell type annotations from the source datasets (i.e. DanioCell and Mokalled Lab database) were compared and verified with our annotations of the integrated dataset.

### Clusterfold similarity test

The “ClusterFold Similarity” package was used to compare the similarities with each cell type across time point (Gonzalez-Velasco et al., 2025). The package quantifies similarity in transcriptional states by comparing gene-level fold-change vectors and thus minimizing batch effects. Similarity tables and matrices were generated in pairwise comparisons between possible combinations of the datasets. Heatmap was generated using GraphPad Prism.

### Cell cycle scoring

Cell cycle scoring was assayed using Seurat “CellCycleScoring” function. Cell cycle phases were determined based on canonical marker gene expression. Each cell was assigned as “G1”, “S”, or “G2/M” phase. Non-dividing cells were defined as those in the G1 phase; and dividing cells were defined as those in the S and G2/M phases.

### Immune and neuronal subset analysis

Clusters comprised of either immune cells or neurons in the integrated dataset were subclustered using the “subset” function for analysis. Each replicate of the subset was again normalized and scaled using “SCTransform” function with glmGamPoi method (Ahlmann-Eltze and Huber, 2021). 50 principal components were used. Further downstream analysis was done as described above for the integrated analysis. For immune subset, resolution = 0.2 was chosen for clustering. For neuronal subset, resolution = 0.9 was chosen for clustering.

### Subset analysis of *sox2*^+^ cells

*sox2*^+^ cells with feature >0 in the “SCT” assay were subclustered using the “subset” function for analysis. Normalization, integration, and downstream analyses were performed using the standard methods described above. Resolution = 0.7 was chosen for clustering.

### Gene Ontology using Metascape

Gene ontology analysis was performed using Metascape. Input and analysis species were set as *D. rerio*. Express analysis was performed for gene ontology. Metascape identified statistically enriched terms (including GO biological processes, Reactome gene set and KEGG pathway) and calculated hypergeometric p-values and enrichment factors. Significant terms were hierarchically clustered into a tree based on Kappa-statistical similarities among their gene memberships. A kappa score of 0.3 was applied to cast the tree into term clusters. The most enriched term in each cluster was chosen as the representative term.

### D-V axis and maturating scoring

The hdWGCNA package was used to identify gene modules with high co-expression coefficient (Morabito et al., 2023). To calculate correlations, Larval EpGlia were assigned a value on a continuous integer scale (e.g. 0, 1, 2) based on their cell type and D-V position. Modules with high correlation to either D-V axis (> |+/- 0.4|) or biological age (> |+/- 0.3|) were selected to establish our scoring framework for Larval EpGlia. The pseudocharacters (D-V or age) were calculated by performing PCA for the eigengene value of select gene modules and using the PC1 as the pseudocharacter value. Adult cells were mapped onto the pseudocharacter axes by projecting cells into the PCA space. With the co-expression network and gene expression matrix set up from larvae, scalar multiplication was performed on the adult EpGlia dataset. The scores for D-V axis and maturation were graphed using “FeaturePlot” and “VlnPlot”.

### Zebrafish and spinal cord transection

Adult zebrafish of the Ekkwill and AB strains were maintained at the Washington University Zebrafish Consortium Facility. All animal experiments were performed in compliance with institutional animal protocols. Fish stocks were tracked and catalogued using VMS-Fish software (Muraleedharan Saraswathy and Mokalled, 2025). Adult male and female animals of ∼2 cm in length (4-6 months of age) were used. Zebrafish were anaesthetized in 0.2 g/l of MS-222 buffered to pH 7.0. Fine scissors were used to make a small incision that transects the spinal cord 4 mm caudal to the brainstem region. Complete transection was visually confirmed at the time of surgery. Injured animals were also assessed at 2 dpi to confirm loss of swim capacity post-surgery.

### Histology

Transverse cryosections of 16 µm thickness were generated from paraformaldehyde-fixed SC tissues. Tissue sections were imaged using a Zeiss LSM 800 confocal microscope or a Zeiss Axioscan.Z1 slide scanner.

For immunohistochemistry, tissue sections were hydrated in PBT then blocked with 5% goat serum for 1 h at room temperature. Sections were incubated overnight at 4 °C with primary antibodies diluted in blocking agent, then washed in PBT, and treated for 1 h in secondary antibodies diluted in blocking agent at room temperature. Primary antibodies used in this study were: chicken anti-GFP (Aves Labs, GFP-1020; 1:1000). Secondary antibodies used in this study were: Alexa Fluor 488, Alexa Fluor 594, and Alexa 647 goat anti-rabbit or anti-mouse antibodies (Jackson ImmunoResearch, 1:200, 103-545-155, 111-605-003, 111-585-003, 115-605-003 and 115-585-003). Hoechst staining was performed according to the manufacturer’s recommendation (Thermo Fisher Scientific, H3570).

The HCR RNA *in situ* hybridization protocol was adapted from Molecular Instruments. Briefly, tissue sections were hydrated and pre-treated either with 0.2% TritonX-100 for 5 min or boiled in Citrate Buffer (10 mM Citric Acid, 0.05% Tween-20, pH 6.0) for 10 min. For blocking, sections were incubated in Hybridization buffer (Molecular Instruments) for 1 h at 37 °C, then incubated in pre-warmed sets of DNA probes diluted to 0.0015 pmol/uL in Hybridization buffer for 48 h at 37 °C. Washes were then performed with Wash Buffer (Molecular Instruments) at 37 °C followed by 5x SSCT (3 M NaCl, 0.3 M Sodium Citrate, 0.1% Tween-20, pH 7.0). For signal amplification, sections were incubated in Amplification buffer (Molecular Instruments) for 1 h at room temperature. Prior to amplification, h1 and h2 hairpins were snap-cooled and diluted 1:50 in Amplification buffer. Amplification proceeded overnight at room temperature in the dark. Samples were then washed in 5x SSCT, 5x SSC and PBT before proceeding to immunohistochemistry. Previously published HCR RNA probes were used for *ctgfa* and *mstnb* (Saraswathy et al., 2022; Zhou et al., 2023).

