## Supplemental Figures for "Spinal cord regeneration deploys adult molecular programs that do not recapitulate embryonic development"

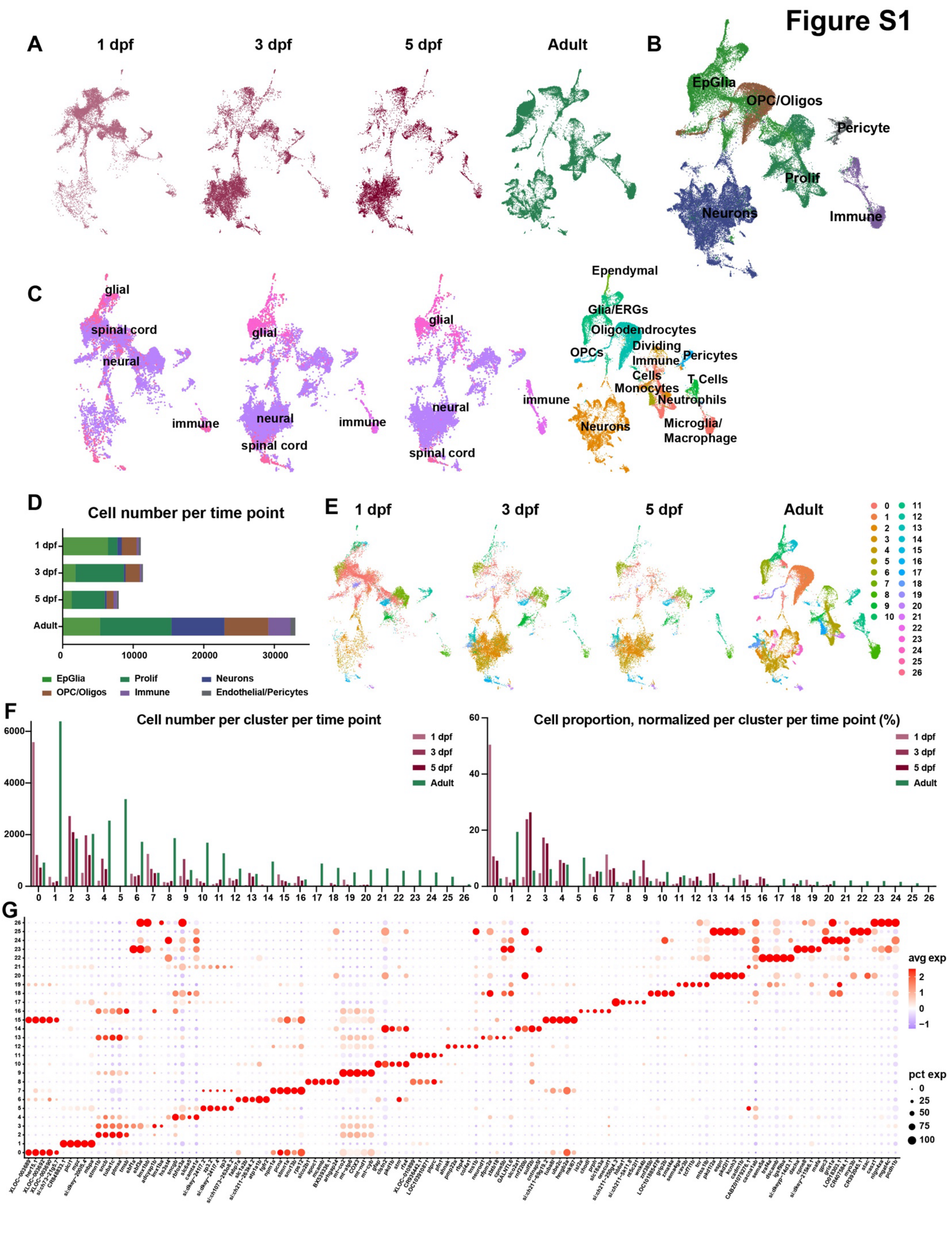
Figure S1
A
1 dpf
3 dpf
5 dpf
Adult
B
EpGlia
OPC/Oligos
Pericyte
Prolif
Neurons
Immune
C
glial
spinal cord
neural
immune
Ependymal
Glia/ERGs
Oligodendrocytes
OPCs
Dividing
Immune
Cells
Pericytes
T Cells
Monocytes
Neutrophils
Neurons
Microglia/
Macrophage
D
Cell number per time point
EpGlia
Prolif
Neurons
OPC/Oligos
Immune
Endothelial/Pericytes
E
F
Cell number per cluster per time point
Cell proportion, normalized per cluster per time point (%)
G
avg exp
pct exp

**Figure S1. Assembly of a single-cell atlas for SC development and adult homeostasis reveals coarse transcriptional differences. (A)** Split UMAP representation of the integrated larval-adult dataset, colored by time point. **(B)** Combined UMAP of the integrated larval-adult dataset, colored by cluster identity. **(C)** Split UMAP representation of the integrated larval-adult dataset, colored by their original annotations in DanioCell (Sur *et al.*, 2023) and in adult SCs (Saraswathy *et al.*, 2024). **(D)** Cell type composition at 1, 3, 5 and 120 dpf. The numbers of cells analyzed for each time point and coarse cluster are shown. **(E)** Split UMAP representation of the integrated larval-adult dataset, colored by cluster identity. **(F)** Numbers (left) and proportions (right) of cells within each cluster from the integrated larval-adult dataset. **(G)** Dot plot shows the top 5 markers for each cluster from the integrated larval-adult dataset. Dot size and color represent percent expressing cells and average gene expression within each cell type, respectively.

Figure S2

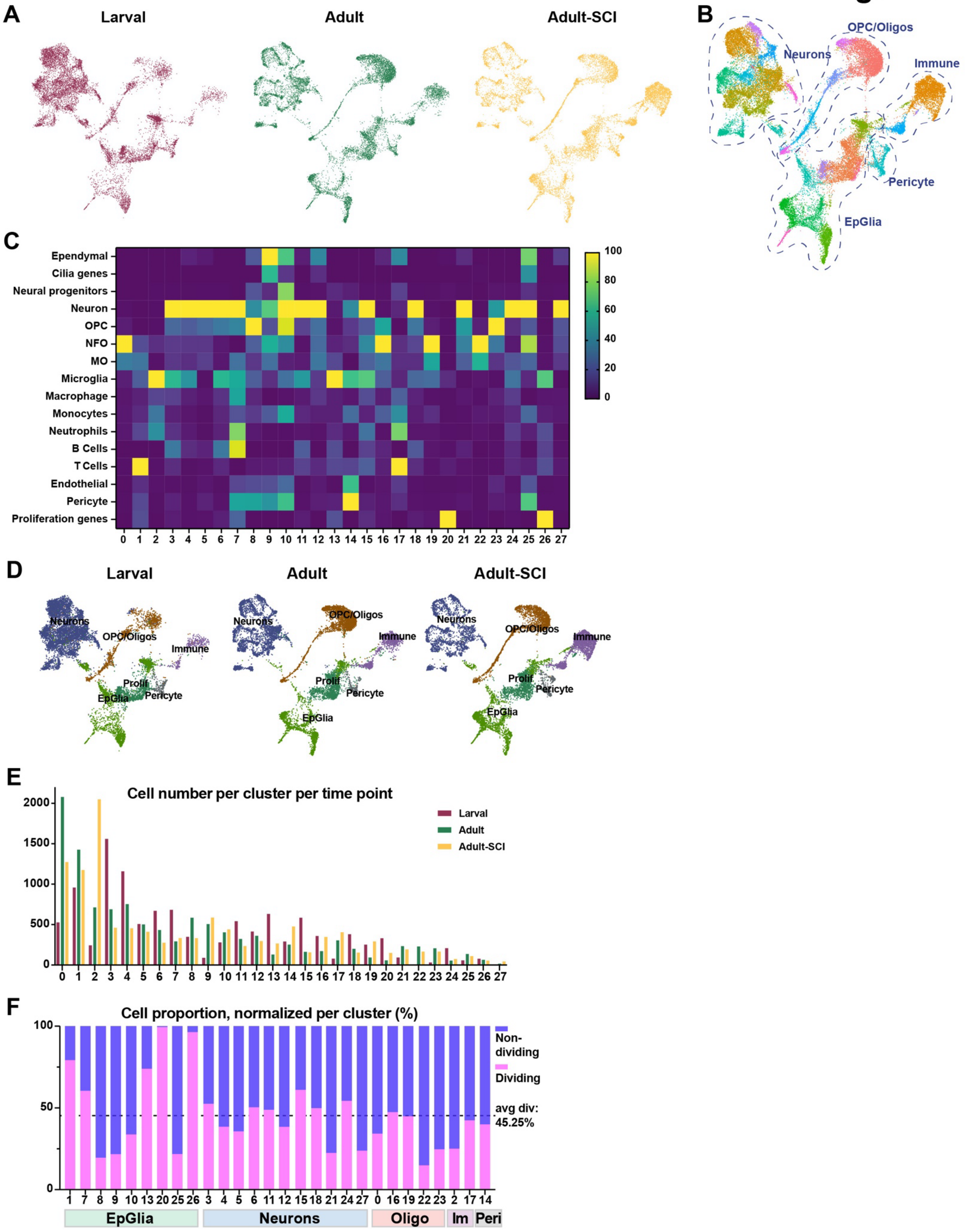
A
Larval
Adult
Adult-SCI
B
OPC/Oligos
Neurons
Immune
Pericyte
EpGlia
C
Ependymal
Cilia genes
Neural progenitors
Neuron
OPC
NFO
MO
Microglia
Macrophage
Monocytes
Neutrophils
B Cells
T Cells
Endothelial
Pericyte
Proliferation genes
0 1 2 3 4 5 6 7 8 9 10 11 12 13 14 15 16 17 18 19 20 21 22 23 24 25 26 27
100
80
60
40
20
0
D
Larval
Adult
Adult-SCI
Neurons
OPC/Oligos
Immune
Prolif
EpGlia
Pericyte
E
Cell number per cluster per time point
Larval
Adult
Adult-SCI
2000
1500
1000
500
0
0 1 2 3 4 5 6 7 8 9 10 11 12 13 14 15 16 17 18 19 20 21 22 23 24 25 26 27
F
Cell proportion, normalized per cluster (%)
100
50
0
Non-dividing
Dividing
avg div: 45.25%
1 7 8 9 10 13 20 25 26 3 4 5 6 11 12 15 18 21 24 27 0 16 19 22 23 2 17 14
EpGlia
Neurons
Oligo
Im
Peri

**Figure S2. Assembly of a single-cell atlas for SC development, adult homeostasis and adult regeneration. (A)** Split UMAP representation of the integrated dataset shown in 2A. **(B)** Combined UMAP of the integrated dataset colored by cluster identitiy. **(C)** Heatmap of cell type probability prediction. Probability is represented on a gradient of high (yellow) to low (purple). **(D)** Split UMAP representation of the integrated larval-adult analysis, split by time point – 1, 3, 5 dpf and adult. Colors represent the annotated cell type. **(E)** Numbers of cells analyzed within each cluster and time point of the integrated larval-adult dataset. **(F)** Proportions of dividing and non-dividing cells within each cluster of the integrated larval-adult dataset.

### Figure S3

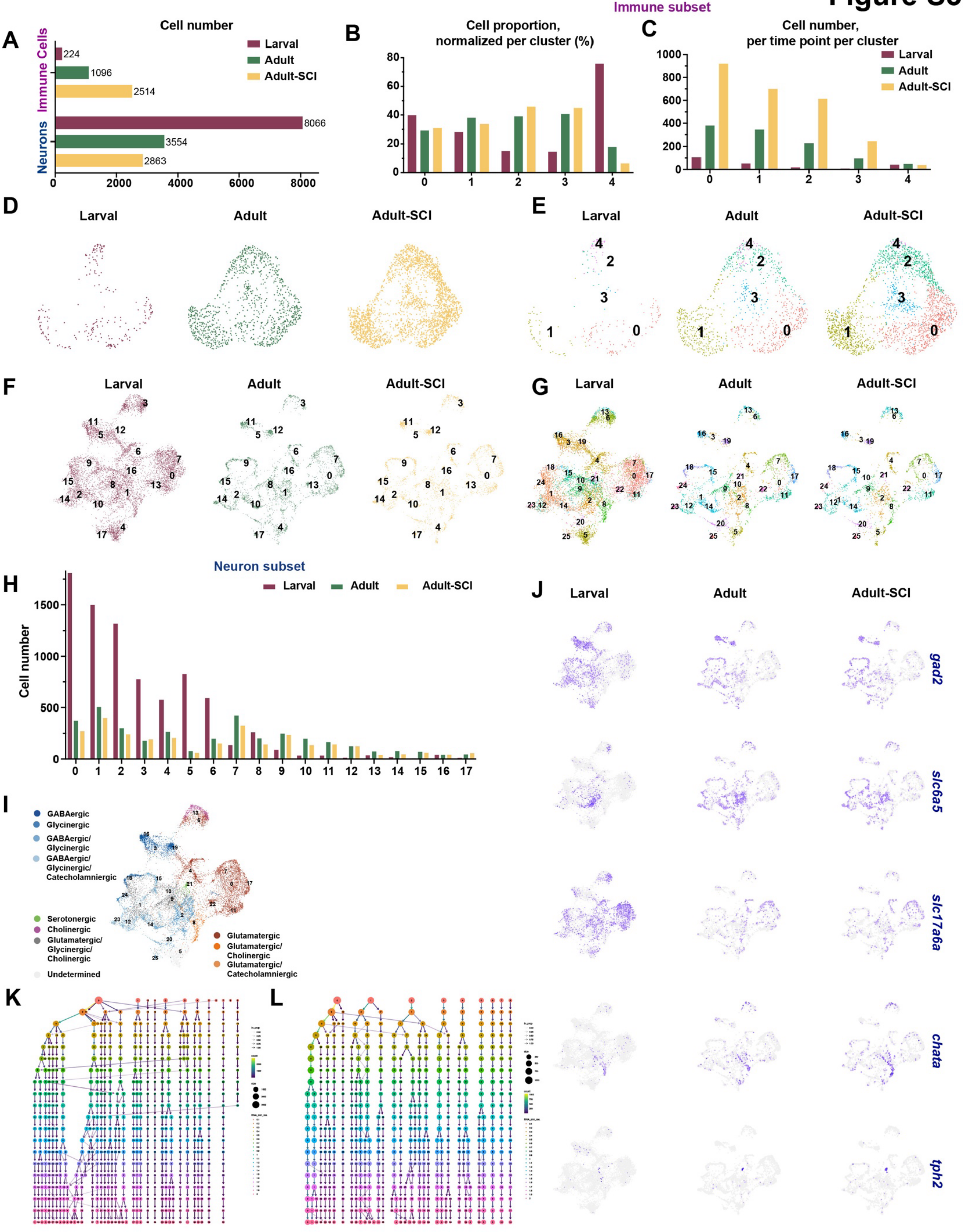
Immune subset
A
Cell number
Immune Cells
Neurons
Larval
Adult
Adult-SCI
224
1096
2514
8066
3554
2863
0
2000
4000
6000
8000
B
Cell proportion, normalized per cluster (%)
C
Cell number, per time point per cluster
D
Larval
Adult
Adult-SCI
E
F
G
Neuron subset
H
Cell number
I
GABAergic
Glycinergic
GABAergic/ Glycinergic
GABAergic/ Glycinergic/ Catecholamniergic
Serotonergic
Cholinergic
Glutamatergic/ Glycinergic/ Cholinergic
Undetermined
Glutamatergic
Glutamatergic/ Cholinergic
Glutamatergic/ Catecholamniergic
J
gad2
slc6a5
slc17a6a
chata
tph2
K
L

**Figure S3. Adult SC regeneration deploys more mature immune cells and increased neuronal diversity compared to development. (A)** Numbers of immune cells and neurons in Larval, Adult and Adul-SCI samples. **(B)** Proportions of immune cells in Larval, Adult and Adul-SCI samples. X axis represent cluster numbers. For each cluster and time point, the numbers of immune cells were normalized to the total number of cells for that cluster. **(C)** Absolute numbers of immune cells per cluster per time point. **(D, E)** Split UMAP representation of immune cells integrated from Larval, Adult and Adul-SCI samples. Cells are color-coded by time point (D), or by cluster identity (E). **(F,G)** Split UMAP representation of neurons integrated from Larval, Adult and Adul-SCI samples. Cells are color-coded by time point (F), or by cluster identity (G). **(H)** Absolute numbers of neurons per cluster per time point. **(I)** Combined UMAP of neuronal clusters, color-coded based on the neurotransmitter properties of each cluster. **(J)** Feature plots for the neuronal markers *gad2* (GABAergic)*, slc6a5* (Glycinergic)*, slc17a6a* (Glutamatergic)*, chata* (Cholinergic) and *tph2* (Serotonergic) in Larval, Adult and Adult-SCI samples. **(K-L)** Clutree shows independent clustering of Larval neurons (K) or Adult neurons (L) at different resolutions (ranging from 0.1 to 2.0).

### Figure S4

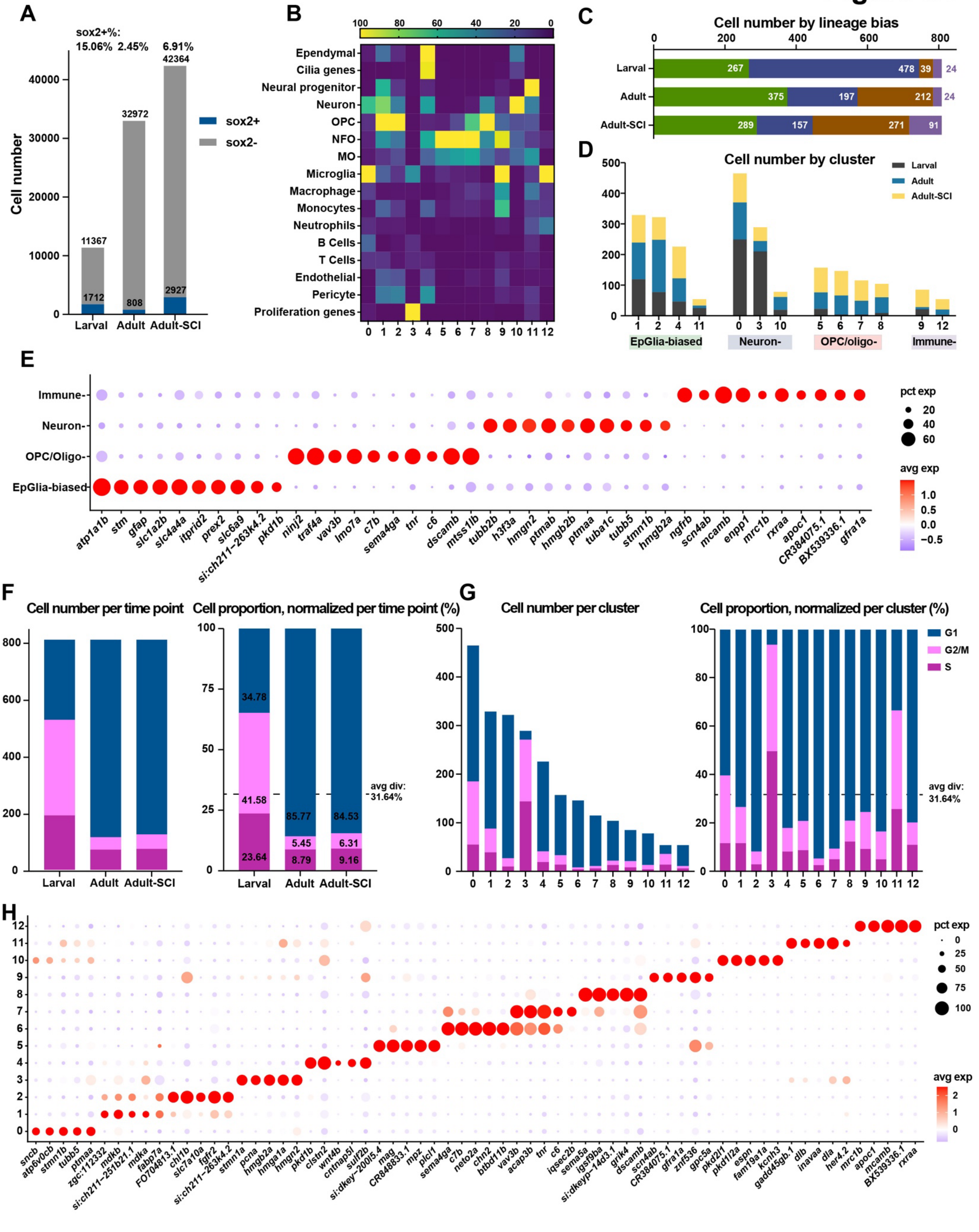
A
sox2+%:
15.06% 2.45% 6.91%
Cell number
sox2+
sox2-
Larval Adult Adult-SCI
B
Ependymal
Cilia genes
Neural progenitor
Neuron
OPC
NFO
MO
Microglia
Macrophage
Monocytes
Neutrophils
B Cells
T Cells
Endothelial
Pericyte
Proliferation genes
C
Cell number by lineage bias
D
Cell number by cluster
Larval
Adult
Adult-SCI
EpGlia-biased
Neuron-
OPC/oligo-
Immune-
E
Immune-
Neuron-
OPC/Oligo-
EpGlia-biased
pct exp
avg exp
F
Cell number per time point
Cell proportion, normalized per time point (%)
avg div: 31.64%
G
Cell number per cluster
Cell proportion, normalized per cluster (%)
G1
G2/M
S
H
pct exp
avg exp

**Figure S4. Subclustering *sox2*$^+$ progenitors during SC development and adult regeneration identifies regeneration-specific clusters. (A)** Numbers of *sox2*$^+$ cells (blue) and *sox2*$^-$ cells (grey) from each time point. The proportions of *sox2*$^+$ cells normalized per time point are listed above the bar graph. **(B)** Heatmap of lineage bias probability prediction. Probability is represented on a gradient of high (yellow) to low (purple). **(C)** Numbers of *sox2*$^+$ cells per lineage bias per time point. **(D)** Numbers of *sox2*$^+$ cells per lineage bias per cluster. **(E)** Dot plot of the top 10 markers of each coarse classification of *sox2*$^+$ cells. Dot size and color represent percent expressing cells and average gene expression within each cell type, respectively. **(F,G)** Numbers and proportions of dividing and non-dividing *sox2*$^+$ cells. The data is represented either per time point (F) or per cluster (G). **(H)** Dot plot of the top 10 markers of each *sox2*$^+$ cell cluster. Dot size and color represent percent expressing cells and average gene expression within each cell type, respectively.

### Figure S5

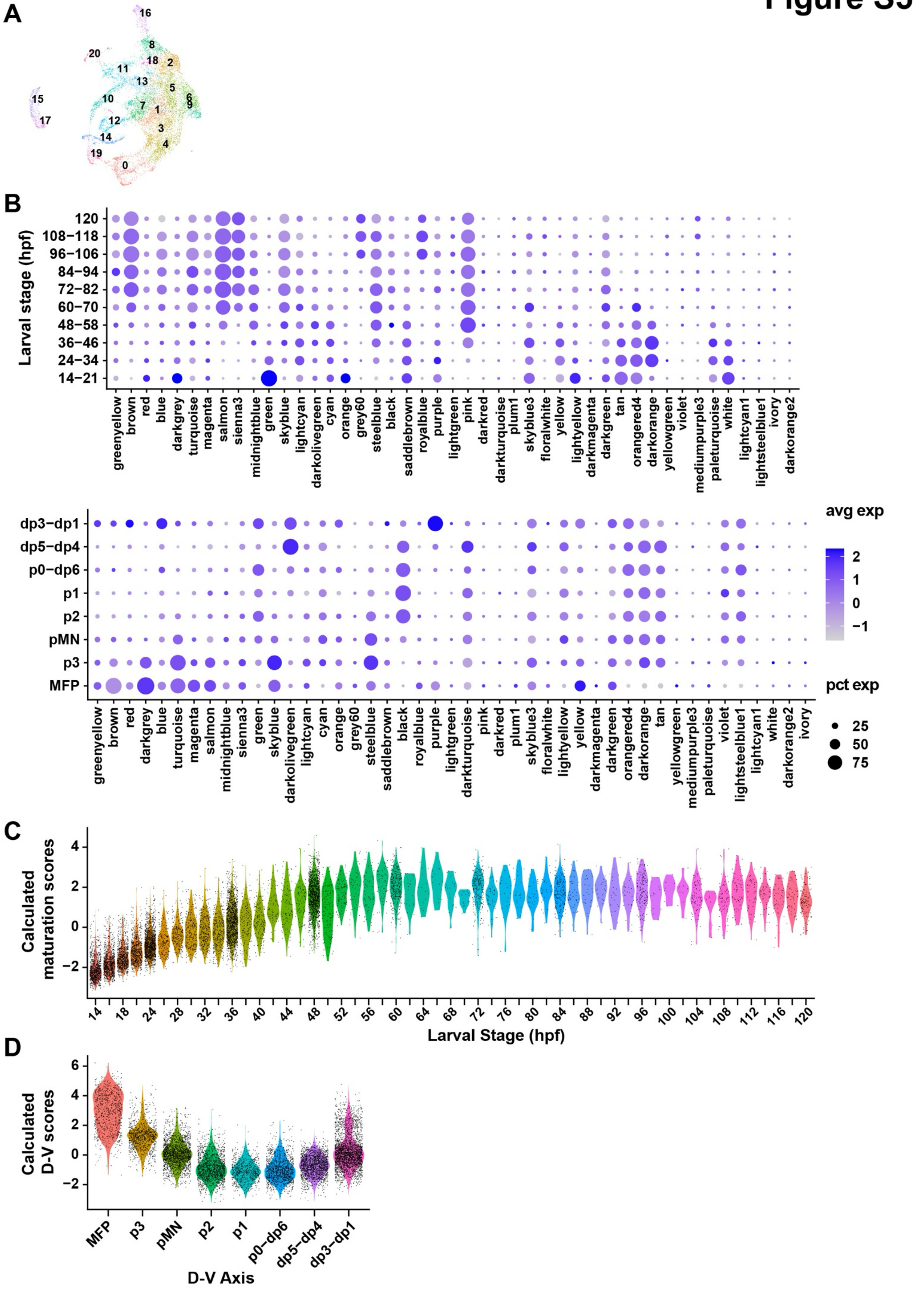

A
16
20
8
18
2
11
13
5
6
9
15
10
7
1
17
12
3
14
4
19
0
B
Larval stage (hpf)
120
108–118
96–106
84–94
72–82
60–70
48–58
36–46
24–34
14–21
greenyellow
brown
red
blue
darkgrey
turquoise
magenta
salmon
sienna3
midnightblue
green
skyblue
lightcyan
darkolivegreen
cyan
orange
grey60
steelblue
black
saddlebrown
royalblue
purple
lightgreen
pink
darkred
darkturquoise
plum1
skyblue3
floralwhite
yellow
lightyellow
darkmagenta
darkgreen
tan
orangered4
darkorange
yellowgreen
violet
mediumpurple3
paleturquoise
white
lightcyan1
lightsteelblue1
ivory
darkorange2
dp3–dp1
dp5–dp4
p0–dp6
p1
p2
pMN
p3
MFP
avg exp
2
1
0
−1
pct exp
25
50
75
C
Calculated maturation scores
4
2
0
−2
14
18
24
28
32
36
40
44
48
52
56
60
64
68
72
76
80
84
88
92
96
100
104
108
112
116
120
Larval Stage (hpf)
D
Calculated D-V scores
6
4
2
0
−2
MFP
p3
pMN
p2
p1
p0–dp6
dp5–dp4
dp3–dp1
D-V Axis

**Figure S5. Establishing and validating a scoring model for maturation and dorso-ventral identity in larval EpGlia. (A)** Combined UMAP of larval EpGlia progenitors spanning 14 to 120 hpf. Colors represent cluster numbers. **(B)** Dot plots represent the expression of all the gene expression modules, or eigengenes, identified in in larval EpGlia at different developmental time points (left) or in D-V location (right). Eigengenes were named after colors and shown on the X axis. Developmental time points (hpf) and D-V axis location are shown on the Y axis. Only select eigenegenes that correlate with either maturation or D-V identity were selected for maturation and D-V scoring and shown in Fig. 5D. **(C, D)** Violin plots validate the calculated maturation scores (C) and calculated D-V scores (D) correlate with the larval stage and positional identities of the samples they are collected from. Larval EpGlia are shown in this validation.
